# Systematic analysis of the molecular and biophysical properties of key DNA damage response factors

**DOI:** 10.1101/2022.06.09.495359

**Authors:** Joshua R. Heyza, Maria Mikhova, Aastha Bahl, David Broadbent, Jens C. Schmidt

## Abstract

Repair of DNA double strand breaks (DSBs) is integral to preserving genomic integrity. Therefore, defining the mechanisms underlying DSB repair will enhance our understanding of how defects in these pathways contribute to human disease and could lead to the discovery of new approaches for therapeutic intervention. Here, we established a panel of HaloTagged DNA damage response factors in U2OS cells which enables concentration-dependent protein labeling. Genomic insertion of the HaloTag at the endogenous loci of the repair factors preserves expression levels and proteins retain proper subcellular localization, foci-forming ability, and functionally support DSB repair. We systematically analyzed total cellular protein abundance, measured recruitment kinetics to DSBs, and defined the diffusion dynamics and chromatin binding by live-cell single-molecule imaging. Our work demonstrates that the Shieldin complex, a critical factor in end joining, does not exist in a preassembled state and Shieldin components are recruited to DSBs with different kinetics. Additionally, live-cell single-molecule imaging revealed the constitutive interaction between MDC1 and chromatin mediated by the PST repeat domain of MDC1. Altogether, our studies demonstrate the utility of single-molecule imaging to provide mechanistic insights into DNA repair, which will serve as a powerful resource for characterizing the biophysical properties of DNA repair factors in living cells.

## INTRODUCTION

Genomic DNA is constantly exposed to a variety of endogenous and exogenous agents that induce DNA double strand breaks (DSBs). These agents include reactive oxygen species, metabolic byproducts, and environmental carcinogens. If left unrepaired, DSBs can lead to the loss of genetic information, chromosome rearrangements, mutations, and telomere fusions that can result in the development of a wide variety of human diseases including cancer, immunodeficiencies, neurological syndromes, and premature aging disorders. In mammalian cells two main pathways exist for resolving DNA DSBs, non-homologous end joining (NHEJ) homologous recombination repair (HR) (Scully et al., 2019). In addition, cells have evolved a complex signaling network that senses DNA breaks, called the DNA damage response (DDR). HR requires a sister chromatid to serve as a template for error-free DSB repair and is limited to S and G2 phases of the cell cycle. In contrast, NHEJ functions independently of a repair template by ligating together broken DNA ends often leading to genomic insertions and deletions. Because of the toxicity of persistent, unrepaired DNA DSBs, NHEJ functions ubiquitously throughout G1, S, and G2 phases of the cell cycle. While cells balance the use of NHEJ and HR particularly in S and G2 phase, NHEJ-dependent repair represents the main pathway resolving the bulk of DSBs arising in human cells either through rapid recruitment of the core NHEJ complex (DNA-PKcs, KU70, KU80, XLF, XRCC4, and LIG4) at unresected DNA ends or after 53BP1-Shieldin mediated end fill-in of previously resected DSBs (Setiaputra & Durocher, 2019).

The DDR is critical for regulating whether DSBs are repaired via HR or NHEJ. DDR action encompasses three specific steps: DNA break detection, DDR signal amplification, and recruitment of processing enzymes and repair effectors. DSB detection is mainly carried out by ATM kinase and the MRN (Mre11, Rad50, and NBS1) complex which associate with DNA breaks leading to phosphorylation of H2AX at S139 (γH2AX) by ATM (Ciccia & Elledge, 2010). Next, DDR signal amplification is mediated by MDC1, RNF8, and RNF168 (Doil et al., 2009; Kolas et al., 2007; Lukas et al., 2004). MDC1 directly binds γH2AX through its C-terminal BRCT domains leading to recruitment of RNF8 and K63-linked polyubiquitylation of linker histone H1 (Mailand et al., 2007; Thorslund et al., 2015). RNF8-mediated histone ubiquitylation is critical for subsequent downstream recruitment of DSB repair effector proteins, such as 53BP1 and BRCA1, to the chromatin regions flanking DSBs. K63-linked ubiquitination serves as a docking site for RNF168 which further amplifies the DDR signal via its E3 ubiquitin ligase activity by depositing monoubiquitin marks onto H2A at K13 and K15 (H2AK13ubK15ub) (Mattiroli et al., 2012).

The critical step in repair pathway choice is the recruitment of either BRCA1 to promote HR or 53BP1 to facilitate NHEJ. One key regulator of this choice is RNF169 which binds H2AK13ubK15ub deposited by RNF168 (Kitevski-Leblanc et al., 2017). RNF169 is a negative regulator of 53BP1 accumulation at DSBs and promotes BRCA1 recruitment (An et al., 2018; R. Menon et al., 2019). 53BP1 binds to H2AK15ub in addition to H4K20me2 which serves as the basis for recruitment of 53BP1 to DSBs (Wilson et al., 2016). 53BP1 binding in turn triggers recruitment and assembly of the 53BP1-RIF1-Shieldin complex where 53BP1 associates with RIF1 which in turn binds to SHLD3, REV7, SHLD2 and SHLD1 (Dev et al., 2018; Gupta et al., 2018; Noordermeer et al., 2018) (Fig. 1A). The Shieldin complex directly binds to ssDNA mediated by the OB-fold domains of SHLD2 (Dev et al., 2018; Gao et al., 2018; Noordermeer et al., 2018). One major regulator of DSB repair pathway choice is the extent of end resection at the DNA break. Resection is carried out by several nucleases including CtIP and the MRN complex, Exo1, and DNA2 (Zhao et al., 2020). Extensive end resection is required for HR while limited resection favors NHEJ. One important function of the Shieldin complex is to protect DNA breaks by competing with HR factors for ssDNA binding, preventing further resection. In addition, the Shieldin complex promotes end fill-in by recruiting Polα/Primase via the CST (CTC1, STN1, and TEN1) complex facilitated by an interaction between SHLD1 and CTC1 (Mirman et al., 2022). By preventing resection and promoting end fill-in the Shieldin complex mediates the formation of DNA ends compatible with NHEJ.

**Figure 1.**
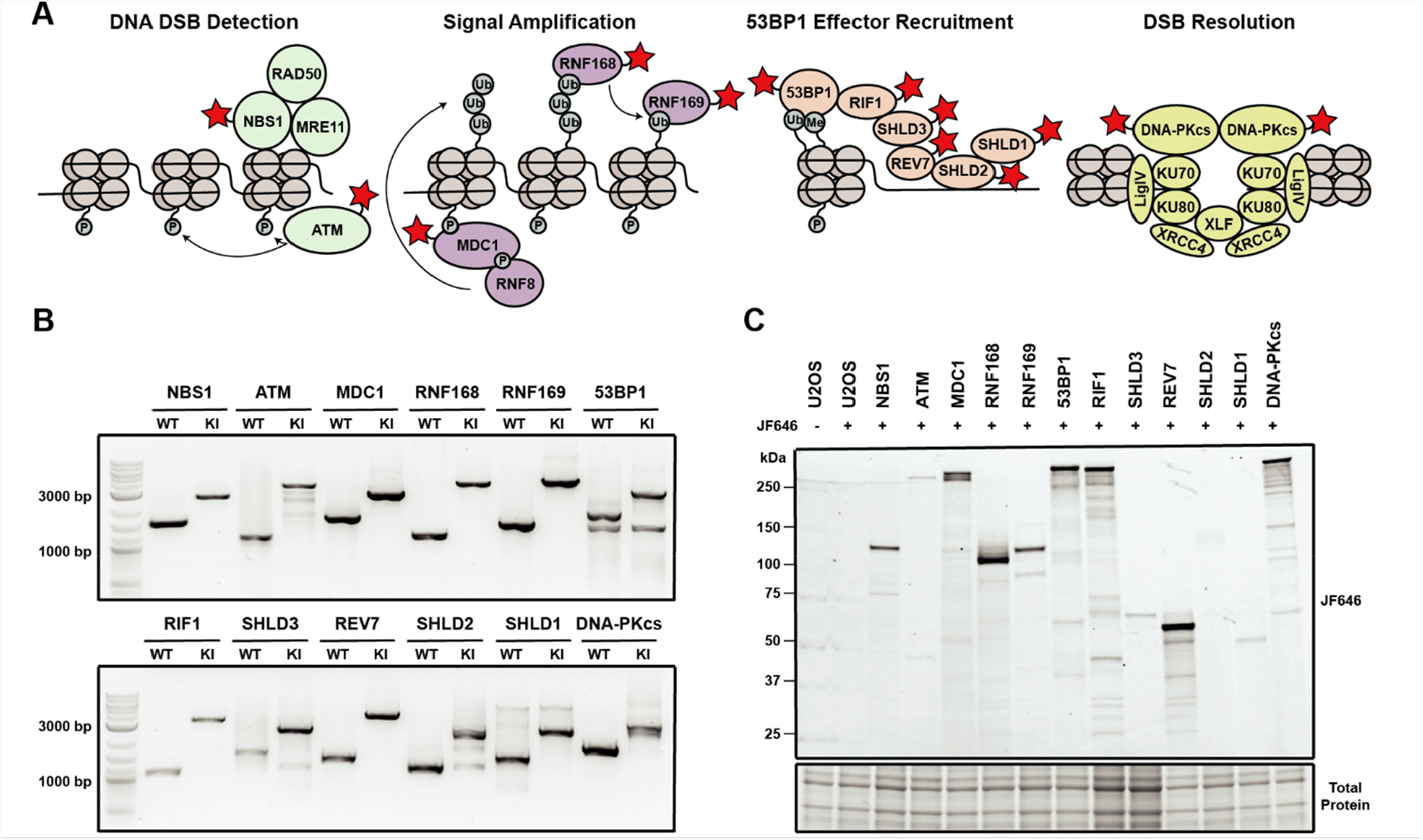
Generation of a panel HaloTagged DNA damage response proteins with CRISPR-Cas9 and homology-directed repair. **A**. Model of the DDR factors HaloTagged by genome editing and their roles in DSB repair. **B**. Agarose gels PCR products amplified from genomic DNA showing insertion of the 3xFLAG-HaloTag into the genomic loci of each tagged DDR factor using primers oriented outside of both left and right homology arms. (WT = wildtype; KI = knock-in) **C**. SDS-PAGE gel showing fluorescently labeled HaloTagged proteins in each cell line after labeling with JF646 HaloTag ligand.

While we have extensive knowledge of the proteins involved in DNA repair, their genetic interactions, and biochemical activities, how their dynamic recruitment to DNA breaks controls repair pathway choice and preserves genome integrity is poorly understood. Importantly, absolute protein abundance, the mechanism by which repair factors search for breaks (e.g. 3D-diffusion, chromatin sampling/scanning), the dynamics of DNA break binding, and the sequence of repair factor recruitment to DSBs are critical determinants of repair pathway choice. In this study we have generated a collection of cell lines that express 12 HaloTagged DNA repair factors from their endogenous genomic loci. The tagged proteins encompass a variety of functions including DSB detection (ATM and NBS1), DDR signal amplification (MDC1 and RNF168) and repair effector recruitment (RNF169, 53BP1, RIF1, SHLD3, REV7, SHLD2, SHLD1, and DNA-PKcs). Using this panel of endogenously edited cell lines we systematically determined the absolute protein abundance of each protein, the kinetics of recruitment to laser-induced DSBs, and defined the diffusion dynamics and search mechanism of all factors using live cell single-molecule imaging. These studies provide new insights into the interplay between RNF169 and 53BP1 in DSB repair pathway choice. While RNF169 and 53BP1 are thought to compete for binding to the same ubiquitin mark, we observe distinct recruitment kinetics to laser-induced DSBs, suggesting that RNF169 binds to its docking site with greater affinity or that additional factors contribute to delayed 53BP1 recruitment. Furthermore, live-cell single-molecule imaging of the lowly abundant Shieldin complex components (SHLD1, SHLD2, SHLD3) demonstrate that the Shieldin complex does not exist as a preassembled complex but rather assembles at DNA lesions. Finally, live-cell single-molecule imaging reveals that MDC1 exists in a constitutive chromatin-associated state, which is mediated by MDC1’s large unstructured PST repeat region and is independent of its BRCT domain. Altogether, our work provides new insight into the molecular mechanism of DNA repair in human cells, establishes a new approach to analyze DNA repair factor recruitment to DNA lesions in living cells, and our panel of cell lines expressing HaloTagged DNA repair factors from their endogenous loci will be a powerful resource for the DNA repair field.

## Results

### Generation of a panel of endogenously HaloTagged DDR Proteins by Genome Editing

To investigate the dynamics of DNA repair proteins at the single-molecule level in living cells, we used CRISPR-Cas9 and homology-directed repair to insert a 3x-FLAG-HaloTag at their endogenous genomic loci in U2OS cells (Fig. S1A). We selected a variety of DNA repair factors that encompass various functional steps of DNA repair and the DNA damage response (DDR) including DNA double strand break (DSB) detection (ATM & NBS1), DDR signal amplification (MDC1 & RNF168), inhibition of DNA end resection (53BP1, RNF169, RIF1, SHLD3, REV7, SHLD2, and SHLD1), as well as DSB resolution (DNA-PKcs) (Figure 1A). Proteins were tagged at either the N-Terminus (MDC1, 53BP1, SHLD3, SHLD2, SDHL1, and DNA-PKcs) or the C-Terminus (NBS1, ATM, RNF168, RNF169, RIF1 & REV7) (Fig. S2B).

We generated clonal cell lines for all targeted proteins and confirmed genome editing using genomic PCR and Sanger sequencing. All cell lines were homozygously edited except for Halo-SHLD2 which had one tagged allele and a frameshift mutation in the second allele (Fig. 1B and Fig. S1C). We obtained at least two clones for all knock-ins except for NBS1, 53BP1 and SHLD2 for which we obtained a single knock-in clone expressing only the HaloTagged protein. The presence of HaloTagged protein was validated by detection of fluorescently labeled proteins in an SDS-PAGE gel after labeling with the cell permeable HaloTag ligand JF646 (Fig. 1C). In addition, we confirmed that all cell lines exclusively expressed the tagged protein by western blotting when antibodies for these proteins were commercially available, which is critical to assess whether the HaloTagged proteins are fully functional (Fig. S1D). HaloTagged proteins validated by Western blotting were expressed at or near endogenous protein levels (Fig. S1D). Despite the commercial availability of antibodies to SHLD2, SHLD1, and RNF169, we were unsuccessful at detecting these proteins at endogenous expression levels. As an alternate approach for proteins where commercial antibodies were not available, we confirmed expression of these proteins by western blotting using an antibody directed to the 3X FLAG epitope (RNF169, SHLD3, SHLD2, and SHLD1) (Fig. S1D). In total, we generated a panel of 12 clonal cell lines that exclusively express HaloTagged DDR factors from their endogenous loci.

### Functional validation of HaloTagged DDR proteins

To ensure that the HaloTagged DDR factors were functional, we assessed protein localization, recruitment to DNA damage induced foci, and clonogenic survival assays after treatment with the DSB-inducing drug Zeocin. To confirm HaloTagging did not interfere with proper protein localization, we imaged JF646-labeled HaloTagged proteins in live cells. As previously described, NBS1, MDC1, 53BP1 RIF1, RNF168, RNF169 and DNA-PKcs localized to the nucleus, while SHLD3, SHLD2, SHLD1 and REV7 were found in the cytoplasm and the nucleus (Fig. 2A), confirming that the HaloTag does not impact the proper cellular localization of these proteins (Noordermeer et al., 2018; Wilson et al., 2016). Most of these proteins also appeared to be excluded from nucleoli (Fig. 2A). HaloTagging ATM at the N-terminus led to nuclear exclusion of the protein. To confirm that HaloTagging does not interfere with the recruitment of the tagged proteins to DNA damage sites, we analyzed their sub-cellular distribution by live-cell imaging one hour after inducing DNA DSBs with Zeocin. While very few foci were detected in the absence of DNA damage in any of the cell lines, we observed dramatic increases in DNA damage induced foci formation for all proteins except for ATM and DNA-PKcs (Fig. 2A). This data confirms HaloTagged DDR proteins are capable of being recruited to DNA damage sites induced by Zeocin.

**Figure 2.**
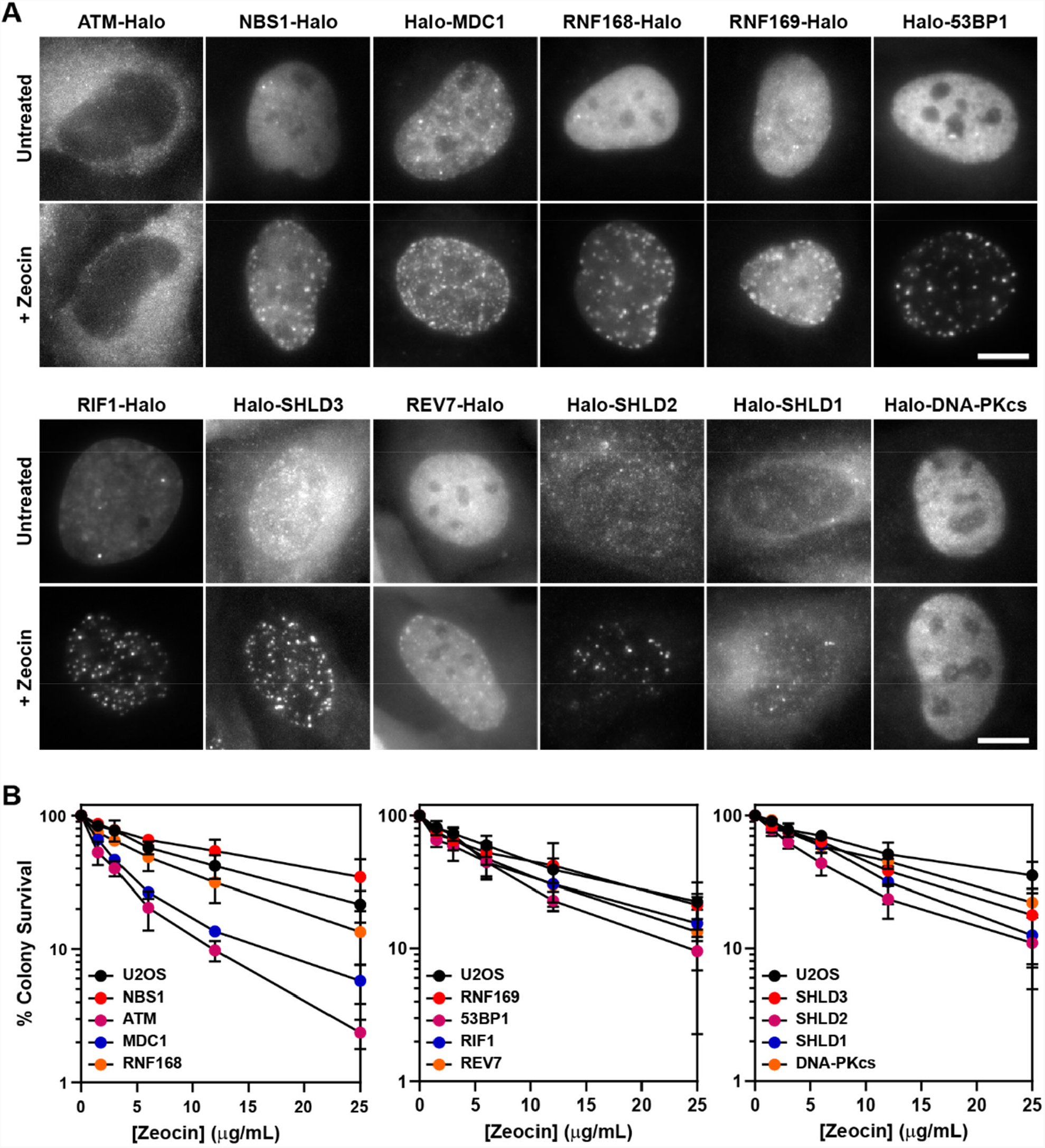
HaloTagged DDR proteins retain proper subcellular localization, foci-forming ability, and are competent for DNA repair. **A**. Representative images of JF646-labeled HaloTagged proteins in the absence or presence of Zeocin in living cells. Data presented show protein cellular localization and foci-forming ability. Scale bar = 10 µm. Images are scaled differently between untreated and treated samples to demonstrate both localization and foci-forming ability. **B**. Clonogenic survival assays representing the Zeocin-sensitivity of each HaloTagged DDR cell line relative to untagged parental U2OS cells. Data presented are the results of at least 3 independent experiments each plated in triplicate ± S.D.

Finally, we tested sensitivity to Zeocin induced DNA damage by clonogenic survival assays of all HaloTagged DDR clones compared to untagged parental U2OS cells. If the HaloTag interfered with protein function in DNA repair, we expected to observe a decrease in clonogenic survival after Zeocin treatment. For most of the HaloTagged proteins Zeocin sensitivity was indistinguishable from the parental U2OS cells, which was consistent between clones (Fig. 2B and Fig. S2A), confirming that the DNA repair function of these proteins is not affected by the HaloTag. Three cell lines were more sensitive to Zeocin than control cells including Halo-ATM, Halo-MDC1, and Halo-53BP1. The HaloTag-ATM cell line was as sensitive to Zeocin as U2OS cells treated with the ATM inhibitor (ATMi) (KU-55933), which could not be alleviated by moving the HaloTag to the C-terminus of ATM, demonstrating that HaloTagging ATM at either end results in a non-functional protein (Fig. S2B, C-D). To assess whether the increased sensitivity to Zeocin induced DNA damage reflected a complete loss of protein function, we established 53BP1 and MDC1 knockout cell lines using CRISPR-Cas9. The Halo-53BP1 and Halo-MDC1 cell lines had modestly increased sensitivity to Zeocin but were significantly more resistant than their knockout counterparts, demonstrating they are partially functional (Fig. S2E, F-G). In summary, with the exception of ATM all of the HaloTagged proteins support repair of Zeocin induced DNA damage, show proper localization within cells, and are robustly recruited to DNA damage induced foci.

### Quantification of absolute cellular protein abundance of DNA repair factors

A key determinant of the kinetics of DDR protein recruitment to DNA damage sites is the absolute cellular concentration of the respective DNA repair factor. For example, the core NHEJ factors, DNA-PKcs, Ku70 and Ku80, are highly abundant proteins which is thought to influence their rapid kinetics of recruitment to DSBs (Carter et al., 1990; Cho et al., 2022; Mimori et al., 1986). Additionally, very little is known about the absolute abundance of members of the Shieldin complex in cells which are thought to be some of the least abundant proteins in the human proteome (Gupta et al., 2018). Thus, determining the absolute cellular protein abundances is critical to establish a mechanistic, quantitative model of the DNA damage response. To measure the protein abundance of the tagged DNA repair factors, we used in-gel fluorescence and flow cytometry-based methodologies. For in-gel fluorescence quantification of the HaloTagged proteins, we modified a method originally described by Cattoglio *et al*. (Cattoglio et al., 2019). To perform these experiments, we generated a standard curve using known quantities of recombinant 3X FLAG-HaloTag labeled with JF646 and cell lysates from a specific number of U2OS cells (Fig. 3A). Two independent clones of each cell line were labeled with 500 nM JF646, which was ∼10x higher than the saturating concentration for the most abundant protein, REV7 (Fig. S3A). Because each protein migrates differently on SDS-PAGE gels based on its size and amino acid composition, we considered the possibility that this may influence the fluorescence of each sample relative to the purified 3X FLAG-HaloTag. To account for differences in the fluorescence signal cause by different migration rates and patterns we cleaved the HaloTag from each protein prior to SDS-PAGE using TEV protease to create an adjustment factor for each protein (Fig. S3B). We obtained a TEV correction factor for each protein except RNF168, RNF169, and MDC1 which all appeared to be degraded after cell lysis which could not be alleviated by including a protease inhibitor cocktail (Fig. S3C). Correction factors ranged 1.04 for SHLD1 to 3.04 for SHLD2 meaning that cleaved HaloTag fluorescence intensity was 1.04x – 3.04x greater for the cleaved HaloTag, than the full-length fusion protein (Fig. S3D).

**Figure 3.**
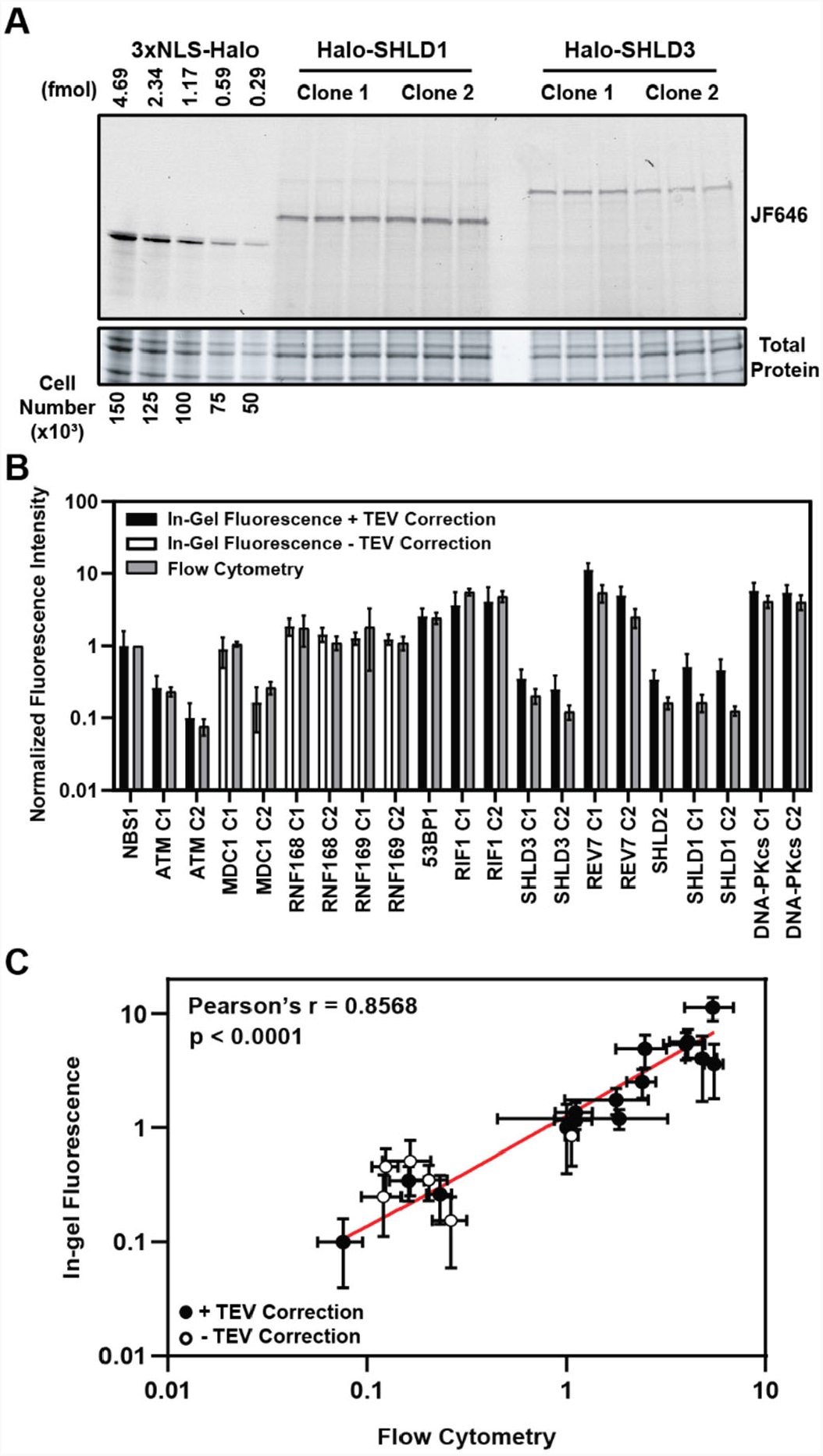
HaloTag enables quantification of absolute cellular protein abundances A. Example image of in-gel fluorescence of JF646-labeled HaloTagged proteins. **B**. Comparison of JF646 fluorescence intensity values (normalized to NBS1-Halo) between in-gel fluorescence after applying the TEV correction factor and flow cytometry. White columns indicate unadjusted samples because of the inability to accurately determine a TEV correction factor due to protein degradation. Data are presented as the mean of all independent experiments ± S.D. **C**. Plot representing the correlation of normalized JF646 fluorescence intensities ± S.D. for each protein between TEV-corrected in-gel fluorescence and flow-cytometry. Data were analyzed by Pearson’s correlation coefficient. White points indicate those proteins for which a TEV-correction factor could not be accurately determined. Red line represents an interpolated standard curve.

After correcting for differences in fluorescence intensity cause by SDS-PAGE migration patterns, HaloTagged protein abundance ranged from ∼1,600 (ATM) – 180,000 (REV7) molecules per cell (Table 1), with ATM (∼1,600 – 4,200 molecules per cell), SHLD1 (∼7,300 – 8,100 molecules per cell), SHLD2 (∼7,300 molecules per cell), and SHLD3 (∼4,000 – 5,600 molecules per cell) clones having the lowest expression level (Table 1). Considering a large proportion of SHLD1, SHLD2, and SHLD3 protein is located in the cytoplasm, the nuclear protein abundance for each of these proteins is even lower which may influence the kinetics of Shieldin complex recruitment in 53BP1-dependent NHEJ. We observed higher protein abundances for factors involved in initial break detection including NBS1 (∼16,000 molecules per cell) and DNA-PKcs (∼86,000 – 91,000 molecules per cell) and those contributing to DDR signal amplification including MDC1 (∼2,400 – 13,000 molecules per cell), RNF168 (∼22,000 – 28,000 molecules per cell), and RNF169 (∼18,000 – 19,000 molecules per cell) (Table 1). The highest protein abundances were observed for 53BP1 (∼40,000 molecules per cell), RIF1 (∼58,000 – 64,000 molecules per cell), REV7 (∼78,000 – 180,000 molecules per cell), and DNA-PKcs (Table 1). Importantly, the differences in absolute protein number between independent genome edited clones could be the consequence of a different number of alleles being modified with the HaloTag. As a complementary approach, we used flow cytometry to measure the fluorescence intensity of JF646-labeled HaloTagged proteins in cells (Fig. S3E). The protein abundance values we obtained by in-gel fluorescence were compared to flow-cytometric quantifications of mean fluorescence intensity for each cell line. Flow-cytometry does not provide an absolute protein number, but instead measures relative protein abundance between the cell lines expressing different HaloTagged proteins. The relative fluorescence levels detected by flow cytometry corresponded well with the relative absolute protein abundance determined by in-gel fluorescence (Fig. 3B,C). Furthermore, results with this complementary approach were comparable to in-gel fluorescence for proteins where a TEV correction factor could not be accurately determined indicating that a TEV correction would have only led to modest changes in overall abundance of these proteins.

**Table 1.** Absolute protein abundances of HaloTagged DNA repair proteins in U2OS cells. Left column: Absolute protein abundance of HaloTagged proteins determined by in-gel fluorescence after adjusting with the TEV correction factor calculated for each protein. An asterisk (*) indicates the three proteins for which an accurate TEV correction factor could not be generated. For TEV-corrected samples, the S.D. includes propogated error. **Right column:** Determination of absolute protein abundances of untagged DNA repair proteins in parental U2OS cells by applying a Western blot correction factor to adjust for differences in expression between HaloTagged and untagged protein. For Western blot corrected samples, S.D. includes propagated error. An asterisk (*) denotes samples where the Western correction was applied to samples that were not TEV corrected. No Data indicates the absence of a commercially available antibody or an antibody that detects endogenous levels of protein expression.

| Protein | HaloTag Protein ± S.D. | + Western Correction |
| --- | --- | --- |
| NBS1 | 15,846 ± 9,547 | 18,055 ± 11,180 |
| ATM C1 | 4,161 ± 1,897 | 4,738 ± 2,379 |
| ATM C2 | 1,578 ± 945 | 3,115 ± 2,365 |
| MDC1 C1 | 13,352 ± 6,001* | 16,030 ± 7,631* |
| MDC1 C2 | 2,434 ± 1,506* | 3,390 ± 2,180* |
| RNF168 C1 | 27,787 ± 7,241* | 4,585 ± 1,819* |
| RNF168 C2 | 21,531 ± 4,801* | 4,678 ± 1,145* |
| RNF169 C1 | 19,013 ± 3,759* | No Data |
| RNF169 C2 | 18,413 ± 3,119* | No Data |
| 53BP1 | 40,162 ± 11,498 | 12,109 ± 5,792 |
| RIF1 C1 | 57,486 ± 28,970 | 9,382 ± 5,765 |
| RIF1 C2 | 64,358 ± 37,504 | 10,398 ± 7,506 |
| SHLD3 C1 | 5,555 ± 1,913 | No Data |
| SHLD3 C2 | 3,953 ± 2,187 | No Data |
| REV7 C1 | 178,131 ± 40,757 | 39,803 ± 10,299 |
| REV7 C2 | 78,340 ± 25,161 | 66,790 ± 42,830 |
| SHLD2 | 7,257 ± 3,104 | No Data |
| SHLD1 C1 | 8,124 ± 4,104 | No Data |
| SHLD1 C2 | 7,257 ± 3,103 | No Data |
| DNA-PKcs C1 | 90,899 ± 26,030 | 115,143 ± 46,933 |
| DNA-PKcs C2 | 85,786 ± 23,508 | 71,488 ± 23,508 |

Finally, using western blotting for each protein where an antibody was commercially available and could detect endogenous levels of protein expression, we quantified the concentration of each HaloTagged protein relative to untagged wild-type protein in parental U2OS cells. Expression of HaloTagged proteins relative to untagged protein in U2OS cells ranged from 0.62x and 0.79x for HaloTagged DNA-PKcs clones to 6.13x and 6.19x for RIF1 clones with 53BP1, RNF168, and one REV7 clone also being overexpressed upon HaloTagging (Fig. S3F). We used the relative expression of each protein compared to untagged protein to calculate the number of molecules per cell for each protein in parental U2OS cells (Table 1). After applying this adjustment factor, DNA-PKcs was the most abundant protein ranging from ∼71,000 – 120,000 molecules per cell. Conversely, ATM and RNF168 had the lowest protein abundances ranging from ∼3,100 – 4,700 molecules per cell. In summary, in-gel fluorescence enabled quantification of absolute protein abundance of HaloTagged proteins in U2OS cells with a dynamic range capable of detecting both lowly and highly expressed proteins.

### Kinetics of recruitment of HaloTag DDR proteins to sites of laser-induced microirradiation

After validating that our panel of HaloTagged DDR factors are functional and proficient in DNA repair, we monitored the kinetics of recruitment for each factor to sites of laser-induced DNA damage. Most previous studies that monitored the kinetics of recruitment of DNA repair factors to laser-induced DNA damage sites have used overexpression of fluorescently tagged proteins, which could influence their recruitment kinetics due to changes in protein levels and competition with the untagged endogenous protein. The use of HaloTagged proteins expressed from their endogenous loci allowed us to more accurately determine the recruitment kinetics of each DNA repair factor to laser-induced DNA lesions. HaloTagged DNA repair factors were labeled with JFX650 and cells were pre-sensitized to laser microirradiation (LMI) with Hoechst (1 µg/mL). LMI reproducibly induced robust protein recruitment for each DNA repair factor (Fig. 4A, Fig. S4A, Movie S1-10). To compare the recruitment kinetics of the DNA repair factors we determined the time to half-maximal accumulation (t_1/2_) (Fig. S4B). DNA-PKcs and NBS1 accumulated rapidly after LMI, with a t_1/2_ = 22.1 s and t_1/2_ = 31.3 s, respectively, consistent with their known roles in the early steps of the DNA damage response (Fig. 4B and Fig. S4B). MDC1 (t_1/2_ = 76.5 s) and RNF168 (t_1/2_ = 69.4 s) arrived at similar times, consistent with rapid ubiquitin deposition at DSBs dependent upon MDC1 mediated RNF8 and RNF168 recruitment (Fig. 4B and Fig. S4B). RNF169 (t_1/2_ = 185.8 s) was significantly delayed compared to RNF168 (Fig. 4B and Fig. S4B), indicating that RNF169 requires chromatin modification by RNF168 prior to its recruitment. 53BP1 and RIF1 both accumulated slowly at LMI sites with comparable kinetics (t_1/2_ = 669.0 s, t_1/2_ = 550.8 s, respectively) (Fig. 4C and Fig. S4B). Surprisingly, REV7 (t_1/2_ = 90.8 s) and SHLD3 (t_1/2_ = ∼1168 s), which are thought to form a complex arrived at LMI induced DNA lesions with distinct kinetics (Fig. 4C and Fig. S4B). In addition, SHLD2 (t_1/2_ = 287.1 s) is recruited more rapidly than 53BP1 and RIF1, which suggests that SHLD2 is not strictly dependent on these proteins and could potentially be recruited to DSBs by its association with ssDNA. We were unable to detect SHLD1 recruitment to LMI induced sites of DNA damage, even though SHLD1 formed DNA damaged induced foci after zeocin treatment. This raises the possibility that SHLD1 recruitment is the key regulatory step in shieldin complex mediated end fill-in. In summary, we have determined the recruitment kinetics of the HaloTagged DNA repair factors expressed at endogenous or near endogenous expression levels. The observed kinetics provide critical insight into the sequence and interdependence of the recruitment of the four shieldin complex components.

**Figure 4.**
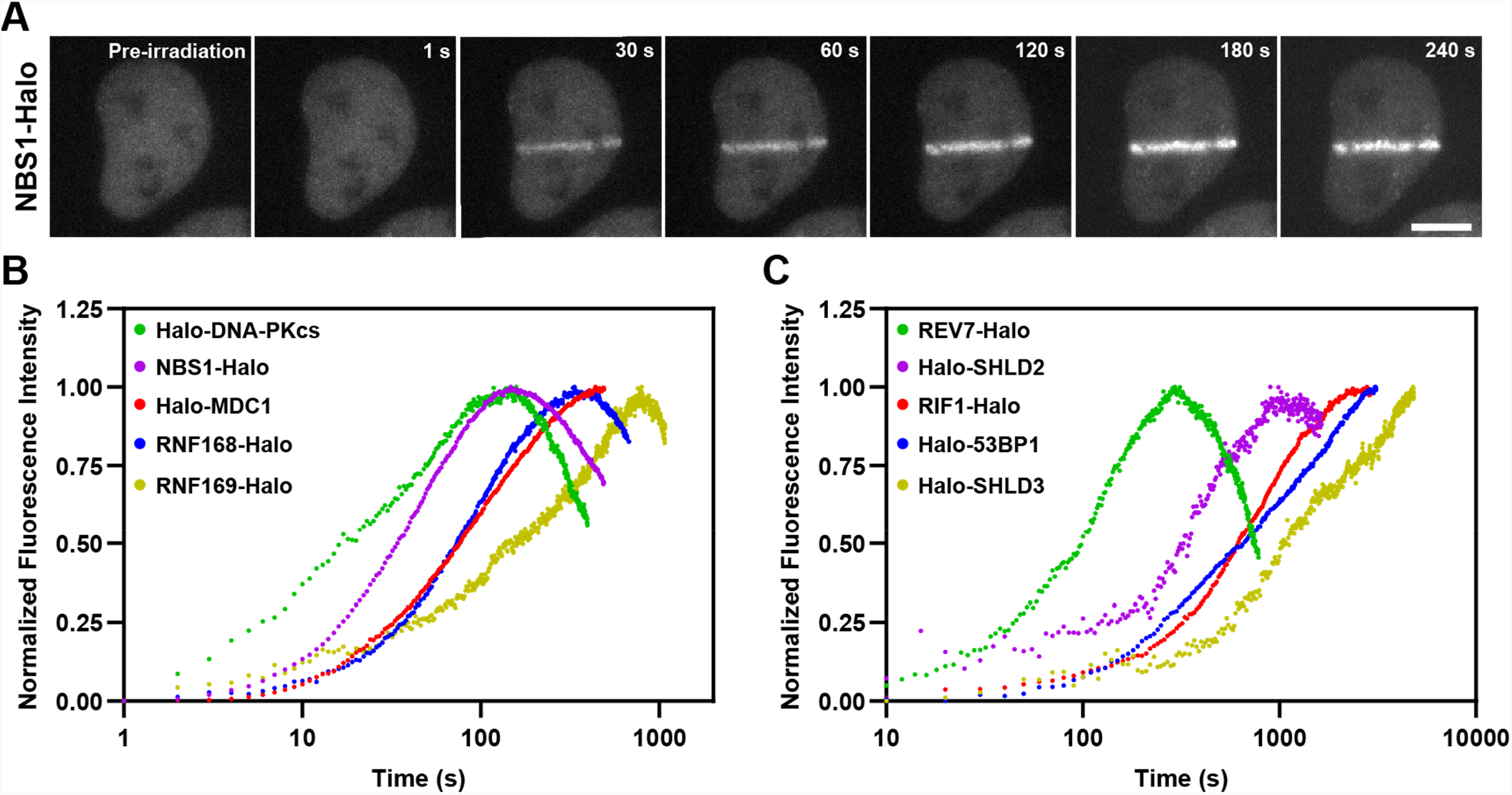
Kinetics of HaloTagged DDR proteins recruitment to sites of laser microirradiation induced DNA breaks. **A**. Representative images of NBS1-Halo (JFX650) recruitment to laser-induced DSBs over time (Scale bar = 10 µm). **B**. Normalized recruitment kinetics of HaloTagged DNA-PKcs, NBS1, MDC1, RNF168, and RNF169 proteins to laser-induced DSBs. **C**. Normalized recruitment kinetics of HaloTagged 53BP1, RIF1, REV7, SHLD2 and SHLD3 proteins to laser-induced DSBs. Data are presented as the average increase in fluorescence post-laser microirradiation normalized to the brightest average frame for each movie. n = 8 - 13 individual cells analyzed for each HaloTag cell line.

### Single-Molecule Live-Cell Imaging of HaloTagged DDR Proteins

The diffusion dynamics of DNA repair factors in the nucleus are a key determinant of their rapid and specific recruitment to DNA lesions. Live-cell single-molecule imaging makes it possible to analyze the diffusion dynamics, chromatin binding, and recruitment to sites of DNA damage of individual DNA repair proteins in their endogenous context. To carry out single-molecule imaging of the HaloTagged DNA repair factors we used a combination of sparse HaloTag labeling and highly inclined laminated optical sheet (HILO) microscopy (Tokunaga et al., 2008), and imaged cells at 138 frames per second to capture rapid diffusion dynamics in both unperturbed conditions and after treatment with Zeocin (Movie S11-S21). Single-particles were automatically detected in each frame, linked into trajectories using the multi-target tracking algorithm (Sergé et al., 2008), and step size distributions were analyzed using the Spot-On tool to extract the diffusion coefficients of freely diffusing (*D*_free_) and chromatin bound molecules (*D*_bound_), as well as the fraction of particles that are associated with chromatin (F_bound_) (Fig. 5A) (Hansen et al., 2018). As model proteins for freely diffusing and chromatin bound factors, we transiently expressed and imaged the HaloTag fused to a 3x Nuclear Localization Sequence (3xNLS) and HaloTagged histone H2B, respectively (Fig. S5A, Movie S22). HaloTag-3xNLS (*D*_free_ = 3.9 µm^2^/s, F_bound_ = 16%) represents the lower bound for static particles and the upper bound for the free diffusion coefficient, while Halo-H2B (F_bound_ = 66%) represents the upper bound for the fraction of bound particles since histone H2B is an integral component of chromatin. For all proteins, we only analyzed nuclear particle trajectories and assumed a two-state diffusion model where particles either freely diffuse or are chromatin bound (Fig. S5B). Importantly, determining *D*_free_, *D*_bound_, and F_bound_ for individual cells (Fig. 5A, Fig. S5C), and combining the step size measurements from all cells from an experimental replicate (Fig. S5D) lead to similar results, indicating that our analysis approach is robust. The diffusion coefficient for freely moving particles ranged from *D*_free_ = 1.0 - 3.7 µm^2^/s, which are all lower than the diffusion coefficient measured for the HaloTag-3xNLS (Fig. 5B). The diffusion coefficients measured for SHLD1 (*D*_free_ = 3.7), SHLD2 (*D*_free_ = 1.8), and SHLD3 (*D*_free_ = 2.3) were significantly different from each other. This suggests that the shieldin complex is not preassembled in the nucleoplasm, which is consistent with the distinct recruitment kinetics to LMI induced sites of DNA damage we observed for the shieldin complex components (Fig. 4C). The diffusion coefficient for bound particles ranged from *D*_bound_ = 0.02 – 0.10 µm^2^/s, which are all faster than Halo-H2B as expected (Fig. S5E). We observed a wide range of the fraction of molecules bound to chromatin (F_bound_ = 23 – 69%) in unperturbed cells (Fig. 5C). Importantly, the chromatin bound fraction of all DNA repair factors analyzed was significantly higher than that of HaloTag-3xNLS, which likely reflects their intrinsic propensity to associate with chromatin (Fig. 5C). Alternatively, the chromatin bound molecules could be the result of low levels of DNA lesions present in control cells. Strikingly, the chromatin bound fraction of MDC1 (F_bound_ = 69%) and RIF1 (F_bound_ = 68%) in unperturbed cells was comparable to that of HaloTag-H2B (F_bound_ = 66%), suggesting that these DNA repair factors are constitutively associated with chromatin and may search for DNA breaks by local chromatin scanning (Fig. 5C).

**Figure 5.**
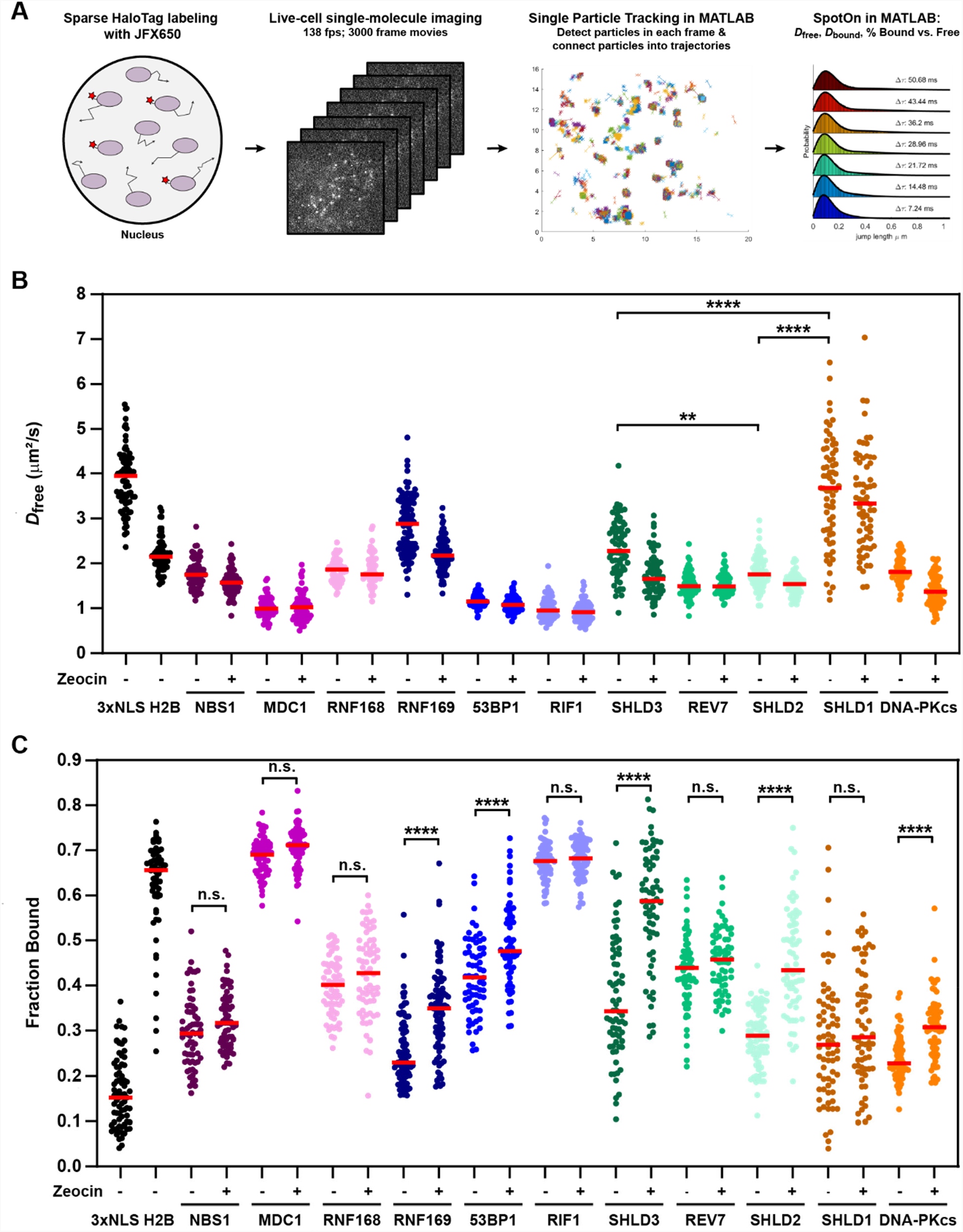
HaloTagged DDR proteins exhibit distinct nuclear diffusion and chromatin binding characteristics. **A**. Graphical representation of the workflow used for live-cell single-molecule imaging of HaloTagged DDR proteins. **B**. Diffusion coefficients for freely diffusing HaloTag DDR proteins present in at least 3 consecutive frames in untreated conditions and post-Zeocin exposure. Values plotted indicate the *D*_free_ for all analyzed tracks per cell with each dot indicating a separate cell that was analyzed. Live-cell single-molecule imaging was performed over 3-4 separate days imaging at least 20 individual cells per condition per experimental replicate (n ≥ 60 cells total for each protein and condition). Red bar = median. **C**. Plot of the Fraction Bound for each HaloTag DDR protein under each condition that were analyzed using a two-state model of diffusion. Each dot represents the fraction bound of each protein in an individual cell (n ≥ 60 cells for each protein and condition). Red bar = median. Data were analyzed by two-way ANOVA with Tukey’s posthoc test. n.s. = not significant. **** = p < 0.0001.

After Zeocin treatment, we observed a significant increase in the fraction of static particles for RNF169, 53BP1, SHLD2, SHLD3, and DNA-PKcs (Fig. 5C), consistent with their recruitment to DNA breaks. Since MDC1 and RIF1 were largely chromatin associated in control cells it is not surprising that we did not observe a further increase in their overall chromatin association after zeocin treatment (Fig. 5C). It is important to note that a change in the fraction bound requires that a significant percentage of the respective DNA repair factor has to be recruited to DNA lesions, which depends on the abundance of the repair factor relative to the number of DSBs induced. NBS1, REV7, and RNF168 are among the most abundant proteins analyzed, which could explain why their F_bound_ did not significantly increase after zeocin treatment (Fig. 5C). Additionally, HaloTagging RNF168 leads to expression ∼4 - 6x that of wild-type protein in U2OS cells which could also contribute to our inability to detect significant changes in protein binding after Zeocin treatment. Despite its low abundance SHLD1 was not recruited to chromatin after DSB induction (Fig. 5C), consistent with its lack of recruitment to LMI induced DNA lesions. Importantly, for all factors that showed an increase in the F_bound_ after zeocin treatment, we observed a decrease in the *D*_free_ (Fig. 5B-C), which is likely the result of a systematic error as a consequence of the global step size fitting in Spot-On. In summary, live-cell single-molecule imaging provides insight into the molecular mechanism by which DNA repair factors search for DNA lesions, reports on complex formation, and allows the analysis of their recruitment to DNA lesions.

### Live-cell single-molecule imaging reveals MDC1’s constitutive chromatin interaction is mediated by its PST repeat domain

Our live-cell single-molecule imaging revealed that MDC1 is constitutively chromatin associated. MDC1 contains two domains that have been implicated in chromatin binding: The BRCT domains that bind to γH2AX (Stucki et al., 2005), and the 13 PST repeats, which have been proposed to associate with the nucleosome acidic patch (Salguero et al., 2019). To dissect the relative contribution of the BRCT and PST repeats domains to the constitutive chromatin association of MDC1, we transiently expressed HaloTagged Halo-MDC1 wildtype (WT), PST deletion (ΔPST), or BRCT deletion (ΔBRCT) mutants in MDC1 knockout (ΔMDC1) cells (Figure 6A-B). We first assessed the ability of these MDC1 variants to localize to DNA damage induced foci. WT MDC1, and MDC1 ΔPST formed foci after zeocin treatment, while MDC1 ΔBRCT was not recruited to DNA damage induced foci (Fig. 6C), consistent with previous observations that BRCT-γH2AX binding is essential for MDC1 foci formation (Stucki et al., 2005). This suggests that the BRCT domains of MDC1 are the primary driver of MDC1 recruitment to DNA lesions. Next, we performed live-cell single-molecule imaging to define the contribution of the BRCT and PST repeat domains to the constitutive chromatin association we observed in unperturbed cells (Fig. 6D, Fig. S6A-B, Movie S23). WT MDC1 transiently expressed in ΔMDC1 behaved identically to endogenously tagged MDC1 (Fig. 6D, Fig. S6A-B, Fig. 5B-C). Deletion of BRCT domains did not alter the diffusion behavior of MDC1 (Fig. 6D, Fig. S6A-B). In contrast, deletion of the PST repeat domain almost completely eliminated the fraction of MDC1 molecules associated with chromatin (Fig. 6D, Fig. S6A-B). This suggests that the PST repeat domain of MDC1 mediates the constitutive chromatin association of MDC1 even in the absence of DNA damage. In addition, the *D*_bound_ of MDC1 was increased upon deletion of the PST repeat domain. This observation indicates that the BRCT domains of MDC1 can rapidly sample histone tails, leading to localized nucleosome scanning without complete dissociation from chromatin. These observations support a model in which MDC1 is constitutively tethered to chromatin by its PST repeat region and is enriched at DNA lesions via the interaction of the BRCT domains with γH2AX histone tails.

**Figure 6.A.**
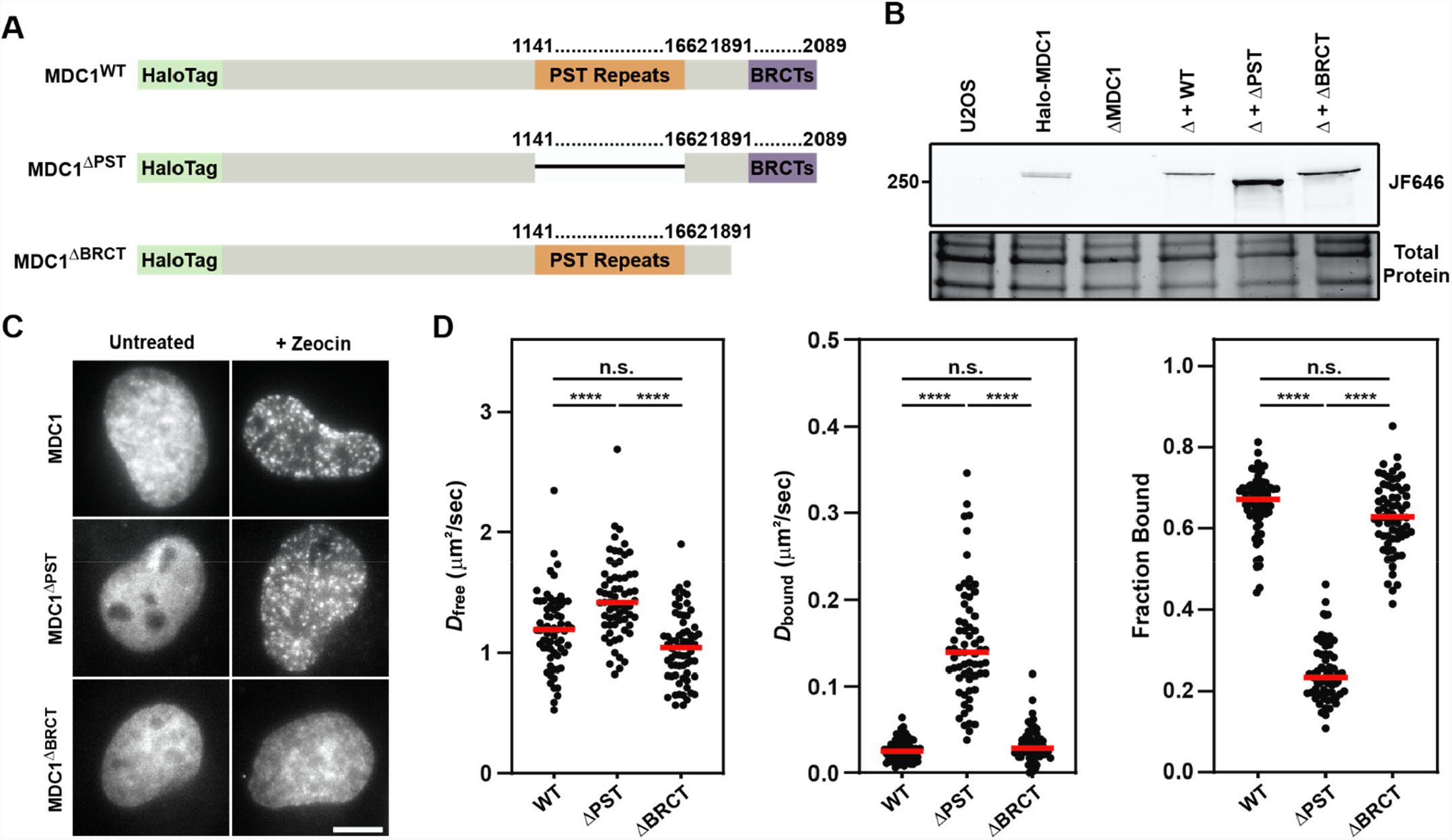
Graphical illustration of the primary sequence of MDC1 indicating the location of the PST repeat and BRCT domain and the associated deletion mutants generated to analyze effects on the MDC1-chromatin interaction. **B**. SDS-PAGE gel of JF646-labeled cells depicting expression of transiently expressed WT, ΔPST, and ΔBRCT MDC1 in Halo-MDC1 knockout cells. **C**. Representative images of transiently expressed, JF646-labeled MDC1 deletion mutants in living cells in the presence of absence of Zeocin. **D**. Results of live-cell single-molecule analysis of transiently overexpressed MDC1 deletion mutants analyzed with single particle tracking and SpotOn. Each dot represents the indicated *D*_free_, *D*_bound_, or Fraction Bound for MDC1 molecules appearing in at least 3 consecutive frames within a single cell. Red bar = median. Data are the combination of all analyzed cells imaged over 3 independent experiments (n ≥ 60 cells total). Data were analyzed by two-way ANOVA with Tukey’s posthoc test. n.s. = not significant. **** = p < 0.0001.

## DISCUSSION

In this study we have developed a panel of cell lines expressing HaloTagged DNA repair factors from their endogenous genomic loci. Using these cell lines we have systematically analyzed the protein abundance, diffusion dynamics, and recruitment to DNA lesions of these factors, which are critical to maintain genome integrity in human cells. Our observations provide new insights into the molecular mechanism and kinetics of the recruitment of the shieldin complex and RNF169 to DNA lesions, which are critical steps in DSB repair pathway choice. In addition, our results reveal how the PST repeat and BRCT domains of MDC1 coordinate its chromatin binding and recruitment to DNA lesions to facilitate DDR signal amplification and DSB repair. In total our work is an important step towards developing a quantitative model of DNA repair in human cells, and provides a number of new tools which will be tremendously useful for the DNA repair research community.

### RNF169 recruitment to DNA lesions precedes 53BP1 to regulate DNA repair pathway choice

Histone tail ubiquitination is a key step in DDR signal amplification and is carried out by the E3 ubiquitin ligases RNF8 and RNF168. The ubiquitination of H2AK13 and H2AK15 by RNF168 is critical for recruitment of BRCA1 and 53BP1 to DSBs. Thus, RNF168 can promote both HR and NHEJ and how effector recruitment downstream of H2AK15 ubiquitination is regulated is a key determinant of repair pathway choice. RNF168 can bind to its own deposited mark to locally amplify the DDR signal (Kitevski-Leblanc et al., 2017). In addition, RNF169 binds to H2AK13K15ub marks, but lacks E3 ubiquitin ligase function. RNF169’s role in DSB repair remains to be fully understood, however it appears to be an important contributor to determining repair pathway choice by competing with 53BP1 for H2AK15ub marks thereby promoting HR, Single-strand annealing, and a-NHEJ (An et al., 2018; R. Menon et al., 2019). Our results demonstrate that RNF168 (t_1/2_ = 69 s) binds to DNA lesions with similar kinetics to MDC1 suggesting that the initial recruitment of RNF8 and the ubiquitination events that lead to RNF168 accumulation occur very fast. In contrast, the downstream recruitment of RNF169 (t_1/2_ = 185 s) and 53BP1 (t_1/2_ = 669.0 s) is delayed substantially relative to RNF168 which places the H2AK15ub mark required for the recruitment of RNF169 and 53BP1 indicating that RNF168-mediated ubiquitination is a slow process or that additional factors are also involved in controlling recruitment of RNF169 to damage sites. Since RNF169 requires the deposition of H2AK15ub by RNF168 for its recruitment its delayed localization to DNA lesions is not surprising. Importantly, RNF169 is recruited well before 53BP1 which effectively reduces the number of H2AK15ub binding sites available for 53BP1 recruitment. Additionally, HaloTagged RNF169 has a protein abundance approximately two-fold lower than 53BP1 which suggests that either RNF169 has a higher affinity for H2AK15ub or that additional events contribute to the delayed recruitment of 53BP1 to DSB sites. Overall, our observations support a model in which RNF169 delays 53BP1 recruitment by competing for H2AK15ub marks, which would allow time for BRCA1 recruitment to occur and end resection to be initiated (An et al., 2017; Poulsen et al., 2012; R. Menon et al., 2019).

### The shieldin complex is recruited to DNA lesions in distinct steps

The shieldin complex is essential to facilitate NHEJ downstream of 53BP1 by recruiting the CST complex and Polα/primase to DSBs. While the pairwise protein and genetic interactions of the shieldin components are well understood, how their dynamic recruitment to DSBs regulates repair via NHEJ was unknown. For instance, it is unclear to what extent shieldin is preassembled either as a complex (SHLD3-REV7-SHLD2-SHLD1), as subcomplexes (e.g., SHLD3-REV7 & SHLD2-SHLD1), or assembled entirely at a DSB. Some evidence supports the hypothesis of assembly at a break. Previous work by others suggests that SHLD2 recruitment is partially independent of SHLD1 while knockdown of REV7 had no impact on the intensity of SHLD3 foci (Noordermeer et al., 2018). It is important to note, that given the high abundance of REV7, knockdown of REV7 might not be sufficient to reveal a loss of function phenotype for this protein. Our live-cell single-molecule imaging demonstrates that SHLD1, SHLD2, and SHLD3 move through the nucleus with distinct diffusion coefficients which strongly suggests that they do not exist in a preassembled state, but rather associate at a DNA lesion.

Quantification of cellular protein abundance for HaloTagged SHLD1, SHLD2, and SHLD3 revealed that these proteins are expressed at comparable low levels. In addition, we observed both cytoplasmic and nuclear localization of each of these proteins by live-cell imaging. Therefore, the nuclear concentration for SHLD1, SHLD2, and SHLD3 is even lower. The low abundance and nuclear exclusion of SHLD1, 2, and 3 suggests that the recruitment of these factors to DSBs is tightly controlled. The low abundance of these proteins also has implications for understanding how these proteins find sites of DNA damage. For example, does recruitment of SHLD1, 2, and 3 occur by freely diffusing molecules encountering DSBs or could enrichment at DSBs be promoted by the establishment of 53BP1-mediated phase separated repair compartments (Kilic et al., 2019). The low abundance of SHLD2 also raises the question of how it can compete for ssDNA binding with RPA which is thought to be more abundant and has a high affinity for ssDNA (Cho et al., 2022; Kim et al., 1992). One explanation could be the local enrichment of SHLD2 at DSBs mediated by 53BP1, comparable to POT1 recruitment to telomeric ssDNA overhangs by the shelterin complex (reviewed in (Litman Flynn et al., 2012)).

Interestingly, we observed large increases in the bound fraction of SHLD2 upon induction of DSBs, while there was no significant increase for SHLD1. This observation would be consistent with two potential models. First, SHLD1 may not bind every break where SHLD2 is present, but rather transiently visit breaks to facilitate Polα-dependent end fill-in via an interaction with CTC1 (Mirman et al., 2022). Another possible explanation is that SHLD1 only binds to one SHLD2 molecule at the 3’ DNA end, while SHLD2 may accumulate either in excess of its substrate (potentially accumulating in 53BP1 condensates), or by forming SHLD2 homopolymers on the ssDNA overhang.

Another unexpected observation arose from monitoring recruitment kinetics of each protein to LMI-induced DSBs. It has been well documented that SHLD3 is the furthest upstream factor in the Shieldin complex coordinating 53BP1-RIF1 with REV7-SHLD2-SHLD1. In experiments overexpressing GFP-SHLD2, knockdown of endogenous RIF1, 53BP1, or SHLD3 led to a marked decrease in GFP-SHLD2 recruitment (Noordermeer et al., 2018). Surprisingly, our data appears to suggest SHLD2 is recruited to LMI DSBs more quickly than RIF1, 53BP1, or SHLD3, having a t_1/2_ of ∼287 seconds before peaking at ∼18 minutes post-irradiation before SHLD3 reaches its t_1/2_. It is important to note that these data are presented based upon normalized fluorescence intensity and therefore does not report on the relative amounts of each protein localized to the laser stripe. A potential explanation for these distinct recruitment kinetics is that at the time that SHLD2 begins to accumulate a sufficient amount of 53BP1, RIF1, SHLD3, and REV7 is already present to recruit SHLD2 to the exposed ssDNA.

Considering that SHLD2 and SHLD3 are present at similar levels in cells this would suggest that substoichiometric amounts of SHLD3 relative to SHLD2 are required to trigger robust SHLD2 localization to DNA breaks. In total, our results suggest a model in which SHLD1, SHLD2, SHLD3, and REV7 are not a preassembled complex but rather associate with each other in the context of a DNA break. SHLD2 can either be recruited by substoichiometric amounts of SHLD3 or independently localize to DSBs via its ssDNA binding activity. Finally, SHLD1 recruitment does not occur as frequently as the other shieldin components and SHLD1 localization to DNA breaks might be a key decision point in repair pathway choice.

### MDC1 and RIF1 are constitutive chromatin-binding proteins

Surprisingly, live-cell single-molecule imaging revealed constitutive chromatin-association of RIF1 and MDC1 in unperturbed cells. The fraction of chromatin bound MDC1 and RIF1 molecules is comparable to what we observe for Halo-H2B. While RIF1 and MDC1 are known to interact with chromatin, the absence of a substantial freely diffusing population leads to several questions for how these proteins function in DSB repair. First, if MDC1 and RIF1 are so immobile, how do they get recruited to DSBs? If the majority of MDC1 and RIF1 are always chromatin-associated, is the small pool of freely diffusing MDC1 and RIF1 sufficient for DSB repair or do chromatin bound molecules need to be locally reorganized or released from chromatin to form a focus? Additionally, RIF1 has many reported nuclear functions that require chromatin binding including telomere maintenance, 3D genome architecture, controlling replication timing, and DSB repair. It is unclear whether the static RIF1 that we observe contributes to one particular function or represents several distinct RIF1 activities. As such, our single-molecule imaging approach would be useful to test how disrupting RIF1 interactions by separation-of-function mutations alters its chromatin binding properties. For MDC1 we demonstrated that the constitutive chromatin association is mediated by its PST repeat domain which was recently reported to associate with the nucleosome acidic patch (Salguero et al., 2019). Some evidence suggests this domain is critical for promoting γH2AX-independent DSB repair, which would suggest that MDC1 constantly scans chromatin searching for DNA damage sites. While our observations would be consistent with that interpretation, it is also possible that MDC1 plays another important as yet unidentified role either in chromatin biology or specifically in DSB repair. Although rarely investigated, MDC1 is annotated as possessing 4 variants produced by alternative splicing, three of which we readily detect in roughly equal amounts. One of these variants has an exclusion of amino acids 1124 – 1410 constituting roughly 50% of the PST repeat region. Because endogenous genome editing preserves this normal splicing pattern, it is possible that this splice variant accounts for the freely diffusing MDC1 that we observe. Future studies will be required to address the role of the PST repeat region and the constitutive chromatin association of MDC1 in DNA repair.

### Using single-molecule live cell imaging to analyze chromatin binding of DNA repair factors

We have developed a new method to analyze the recruitment of DNA repair factors to chromatin and DNA lesions using live cell single-molecule imaging. DNA repair factors that display a significant change in their chromatin bound fraction after DNA damage induction (e.g. DNA-PKcs, SHLD2, SHLD3, RNF169) can be studied using this approach. Importantly, we only tested zeocin induced DSBs in this study. It is possible that other means of inducing DNA lesions (e.g. PARP inhibitors, topoisomerase inhibitors) could result in significant changes in the chromatin bound fraction of other DNA repair factors. Previously, quantitative analysis of the recruitment of DNA repair factors to DNA lesions in living cells was limited to using laser microirradiation (LMI). LMI induces complex lesions, and the precise chemistry of the damaged DNA cannot be controlled. Our single-molecule imaging based approach opens the door to quantitatively analyze the binding of DNA repair factors to DNA damage sites induced with a wide range of agents that lead to chemically well-defined DNA lesions. In addition, single-molecule imaging can directly measure kinetic parameters such as the dissociation rate of DNA repair factors from DNA damage sites, which will significantly advance our quantitative understanding of DNA repair in human cells.

In total, the results described in this study provide important insights into the protein abundance, diffusion dynamics, and recruitment kinetics to DNA breaks of a range of DNA repair factors. Our observations revealed that MDC1 and RIF1 constitutively associate with chromatin and demonstrate that the shieldin complex components are assembled in the context of DNA breaks but otherwise act independently. In addition, the collection of cell lines we have generated and the single-molecule imaging-based analysis of DNA repair factor recruitment to DNA breaks will be valuable tools for scientists studying DNA repair in human cells.

## DATA AVAILABILITY

All plasmids used for genome editing/transient overexpression will be deposited on Addgene. HaloTagged cell lines will be made available upon request.

## ACKNOWLEDGMENTS

We are grateful to Dr. Kathy Meek for providing the DNA-PKcs antibody. We thank Dr. Daniel T. Youmans and Dr. Thomas R. Cech for providing the plasmid for recombinant production of the 3xFLAG-HaloTag protein. We thank Dr. Eric Patrick for contributing to the purification of the 3xFLAG-HaloTag protein.

## Author contributions

Conceptualization: J.R.H. and J.C.S.: Experiments: J.R.H., M.M., A.B., D.B.: Data Analysis: J.R.H., M.M., D.B.: Writing—Original Draft: J.R.H.: Writing—Review and Editing: J.R.H and J.C.S.

## MATERIALS AND METHODS

### Cell Lines and Cell Culture

U2OS cells were cultured in RPMI containing 10% fetal bovine serum (FBS) and 100 units/mL penicillin and 100 µg/mL streptomycin in a humidified incubator maintained at 37º C with 5% CO_2_. For live-cell imaging experiments, cells were plated onto 24-well glass bottom plates and imaging was conducted using CO_2_-independent medium containing 10% FBS, 100 units/mL penicillin and 100 µg/mL streptomycin at 37º C and 5% CO_2_.

### Molecular Cloning, Plasmids and Genome Editing

All gRNAs were cloned into BpiI-digested px330 backbone using standard procedures and ssDNA oligos were purchased from IDT. All homology-directed repair (HDR) donor plasmids were cloned using Gibson Assembly into pFastBac Dual backbone linearized with HpaI. Gibson Assembly for each HDR donor consisted of three inserts including a left homology arm, right homology arm and an intervening sequence containing either the N-terminal or C-terminal HaloTag sequence. The N-terminal tag consists of a 3x FLAG tag, an inverted SV40 promoter and Puromycin resistance cassette (Puro^R^) flanked by LoxP sites (to select for edited cells), the HaloTag, and a TEV protease cleavage site, followed by a short peptide linker. Conversely, the C-Terminal tag consisted of a short peptide linker and TEV protease cleavage site, followed by the HaloTag and the 3X FLAG epitope, with the Puro^R^ cassette oriented 3’ to the 3X FLAG tag. This approach allowed for selectable direct insertion of the HaloTag at the C-Terminus of a protein without the need for the additional step of Cre-Lox recombination to remove the Puro^R^ cassette. Homology arms consisted of >250 bp homologous to the genomic DNA directly upstream and downstream of the Cas9 cleavage site and were either ordered as double-stranded gene fragments from IDT or PCR amplified from genomic DNA. MDC1 deletion mutants were generated using rationally designed PCR primers to amplify MDC1 cDNA. Cloning of the Halo-MDC1 deletion mutants was done by Gibson Assembly into pRK2. All plasmids were confirmed for proper insertion/assembly by Sanger sequencing. Halo-H2B plasmid was kindly provided by Dr. Anders Hansen (Hansen et al., 2017) and the Halo-3xNLS plasmid was previously established and described(Al-Masraf et al., 2021).

Halo-MDC1 deletion mutants were expressed transiently by transfecting ∼ 5 × 10^5^ cells with 1 µg of plasmid DNA using FuGene 6 (Promega). For genome editing, ∼ 5 × 10^5^ U2OS cells were transfected using FuGene 6 with either 500 ng or 1 µg of each gRNA/Cas9 and HDR plasmid in 6 well plates. Approximately 2-3 days post-transfection, edited cells were selected for with puromycin (1 µg/mL). After puromycin selection, cells were allowed to grow for ∼ 2-3 weeks followed by sorting for single-cell clones. N-terminally edited cells were transfected with a plasmid encoding Cre to recombine out the Puro^R^ cassette generating a 3xFLAG-HaloTagged protein. Cells were labeled with JF646-HaloTag ligand (JF646) and sorted based on JF646 signal (Grimm et al., 2015). For knockout of Halo-MDC1 and Halo-53BP1, cells were transfected with two gRNA plasmids. Cells were isolated for single-cell clones by labeling with JF646 and sorting JF646-negative cells. Knockout was confirmed by loss of fluorescence by SDS-PAGE, Western blot, Sanger sequencing and for MDC1 with Inference of CRISPR Edits (ICE) (Conant et al., 2022).

### SDS-PAGE and Western Blot

Mini-PROTEAN TGX stain-free gels (BioRad) were used for SDS-PAGE for most proteins except for MDC1, DNA-PKcs, and RIF1 where homemade 6% polyacrylamide gels were used. Protein samples were made by lysing cells with 2x Laemmli buffer with β-mercaptoethanol. For in-gel fluorescence to detect HaloTagged proteins, cells were labeled with ∼150 - 500 nM JF646 or JFX650 HaloTag ligand (JFX650; similar to JF646 but with improved photostability) for 30 minutes, washed with complete medium three times, allowed to rest for ∼10 minutes, and subsequently washed twice with PBS before being lysed with buffer (Grimm et al., 2021). For in-gel fluorescence measurements of protein abundance, cells were plated in triplicate in 24 well plates, the following day samples were labeled with 500 nM JF646 for ∼30 minutes, washed with PBS three times, lysed in 2x Laemmli buffer with β-mercaptoethanol, and boiled for 5 minutes at 95° C prior to gel loading. Standards were prepared using standards containing known femtomolar concentrations of FPLC-purified, JF646-labeled 3X-FLAG-HaloTag prepared in aliquots containing known cell concentrations, boiled for five minutes, and frozen at -80° C until use. Fluorescence was detected using the Cy5.5 filter on a BioRad Chemidoc. Stain-free detection of protein loading was detected using the Stain-Free filter on a BioRad Chemidoc after 45 second UV activation. For protein abundance, the number of molecules were calculated by comparing the fluorescent signal for each protein relative to the standard curves for cell number and JF646-labeled 3X-FLAG-HaloTag. For TEV corrections, ∼120,000 cells were labeled with 500 nM JF646 for 30 minutes, washed once with PBS and harvested with 5 mM EDTA in PBS. Samples were lysed in 60 µL CHAPS lysis buffer and 20µL incubated on ice with 5 units of TEV protease (New England Biolabs, P8112) for 30 minutes at which time 5 µL 6x SDS sample buffer was added to 20 µL of the digested and undigested sample and boiled at 95° C for five minutes. For TEV correction experiments where protease inhibitors used, samples were pre-treated with 10 µL of protease inhibitor cocktail (Sigma, P8340) for 30 minutes on ice followed by the addition of TEV protease. Comparisons between the JF646 signal ± digestion by TEV protease were performed by comparing the JF646 intensity for each sample after normalizing to stain-free protein signal. Transfers onto PVDF or Nitrocellulose membrane were conducted using either a Trans-Blot Turbo system (with Turbo transfer buffer) (BioRad) or by traditional wet tank transfer using CAPS Buffer with 10% Methanol (for DNA-PKcs, 53BP1, RIF1, and MDC1). Antibodies used were anti-FLAG-HRP (Sigma-Aldrich, A8592, 1:5000 dilution), anti-DNA-PKcs (a gift from Dr. Kathy Meek, 1:1000 dilution), anti-ATM (Santa Cruz, sc-135663, 1:1000), anti-NBS1 (BioRad, VMA00403, 1:1000), anti-MDC1 (Novus Biologicals, NB100-395, 1:1000), anti-RNF168 (GeneTex, GTX129617, 1:1000), anti-53BP1 (Novus Biologicals, NB100-304, 1:1000), anti-RIF1 (Bethyl Laboratories, A300-568A, 1:1000), anti-REV7 (Abcam, ab180579, 1:1000), goat anti-mouse HRP (Invitrogen, 31430, 1:2000), goat anti-rabbit HRP (Invitrogen, 31460, 1:2000).

### Purification of recombinant 3xFLAG-HaloTag protein

The plasmid encoding 6xHIS-3xFLAG-HaloTag protein (a kind gift from Dr. Daniel T. Youmans and Dr. Thomas R. Cech) was transformed into OneShot™ BL21(DE3) cells (Invitrogen). Cultures were grown to an OD600 of 0.6, induced with 1 mM IPTG, and grown at 18 °C for 16 hours shaking at 180 RPM. Bacterial cells were harvested by centrifugation at 5000xg for 15 min at 4 °C and frozen at -80 °C for 1 hour. Frozen cell pellets were resuspended in lysis and wash buffer (50 mM sodium phosphate buffer pH 8.0, 300 mM sodium chloride, 10 mM imidazole, 5 mM beta-mercaptoethanol). Cells were lysed by addition of 0.5 mg/ml lysozyme and sonication (40% amplitude, 90 seconds of sonication, 10 second pulses, 20 second pause, Fisherbrand™ Model 505, 0.5 inch tip) in an ice water bath. Cell lysates were cleared at 40,000xg and 4 °C for 30 minutes and incubated with fast flow nickel Sepharose (Cytiva) for 1 hour at 4 °C. The resin was washed three times with lysis and wash buffer and the 6xHIS-3xFLAG-HaloTag was eluted in elution buffer (50 mM sodium phosphate buffer pH 7.0, 300 mM sodium chloride, 250 mM imidazole, 5 mM beta-mercaptoethanol). The 6xHIS-3xFLAG-HaloTag was further purified using size exclusion chromatography using a superdex 75 column into 50 mM Tris pH 7.5, 150 mM potassium chloride, 1 mM DTT. Peak fractions were combined, concentrated, supplemented with 50% glycerol, snap frozen in liquid nitrogen and stored at -80 °C. To fluorescently label the 6xHIS-3xFLAG-HaloTag, we incubated the protein with a 2-fold excess of JF646 HaloTag-ligand overnight at room temperature. Excess fluorescent dye was removed by size exclusion chromatography using a superdex 75 column into 50 mM Tris pH 7.5, 150 mM potassium chloride, 1 mM DTT. The protein concentration and labeling efficiency was determined by absorption spectroscopy using ε_280nm_ = 41,060 M^-1^ cm^-1^ for the 6xHIS-3xFLAG-HaloTag (calculated using primary protein sequence and the ExPASy ProtParam tool) and ε_646nm_ = 152,000 M^-1^ cm^-1^ for the JF646 fluorescent dye (Grimm et al., 2015).

### Clonogenic Survival Assays

On the day prior to treatment cells were seeded in triplicate at a density of 500 cells per well in six-well plates. The following day, cells were treated with Zeocin (Gibco) at the indicated concentrations for four hours in complete medium. After treatment, medium was removed and replaced with fresh complete medium. For assays including the ATM inhibitor (ATMi) (KU-55933; Selleckchem), cells were pre-treated for 2 hours, during Zeocin treatment, and for 24 hours post-Zeocin with 10 µM ATMi. Cells were allowed to grow until colonies reached a size of >50 cells (approximately 7-10 days) at which time media was removed, wells washed with PBS and cells fixed and stained in crystal violet solution (20% ethanol and 1% w/v crystal violet). After incubation excess staining solution was removed and plates gently washed in diH2O and air-dried. Plates were imaged on a BioRad Chemidoc using the Coomassie Blue filter set. Colony counts were determined using ImageQuant TL 8.2. All colony survival assays were performed in triplicate at least three times and are plotted as the average ± S.D.

### Flow Cytometry

Four independent experiments were performed to quantify relative protein abundance using flow cytometry. On the day prior to performing flow cytometry, ∼100,000 cells of each HaloTagged protein clone were seeded into 24-well plates. The next day, cells were labeled with 500 nM JF646 for ∼30 minutes, washed three times with PBS, and allowed to rest for five minutes in complete medium to allow unbound dye to leak out of the cells. Cells were harvested using 5 mM EDTA in PBS, fixed in 2% paraformaldehyde for 10 minutes, washed once with PBS and resuspended in PBS with 1% bovine serum albumin. Samples were run on a Cytek Aurora spectral flow cytometer and >7,500 events were collected per protein for each replicate to determine mean fluorescence intensity. Data were analyzed in FCS Express 7.

### Laser Microirradiation

Laser microirradiation (LMI) was carried out on a Olympus IX83 inverted microscope equipped with a 4-line cellTIRF illuminator (405 nm, 488 nm, 561 nm, 640 nm lasers), an Excelitas X-Cite TURBO LED light source, a Olympus UAPO 100x TIRF objective (1.49 NA), a CAIRN TwinCam beamsplitter, 2 Andor iXon 897 Ultra EMCCD cameras, a cellFRAP with a 100 mW 405 nm laser, an environmental control enclosure and operated using the Olympus cellSense software. Cells were seeded the day before LMI onto a 24 well glass bottom plate. On the day of the experiment, cells were labeled with 150 nM JFX650 for 30 minutes, washed three times with complete media and presensitized with Hoechst (1 µg/mL) for ten minutes before washing and adding fresh complete medium. Plates were placed on the microscope stage which was pre-warmed to 37ºC with 5% CO_2_. Cells were irradiated using a 20 ms pulse at 25% laser power using drawn interpolated lines or a diffraction limited spot (only used for REV7 due to diffuse staining pattern which made it difficult to quantify over time). Images were acquired using a 100x objective and fluorescence imaged with excitement by the 630 nm LED light source at various rates (e.g., every 1 s, 2 s, 5 s, or 10 s). To quantify the fluorescent signal after LMI, we converted movies to .tif files. Irradiated cells were cropped and images drift corrected using NanoJ in Fiji (Laine et al., 2019). Using drift corrected movies, an ROI was placed around the irradiated area and we measured the mean intensity within the ROI over time. Baseline fluorescence was subtracted from each sample in order to calculate the relative increase in fluorescence that accumulated over time. To normalize fluorescence intensity values, each cell had its highest intensity frame set to one in order to normalize fluorescence intensity values between samples. For averaging normalized intensity values for all cells, the frame number with the highest average intensity was set to one. Data were plotted in GraphPad Prism and recruitment half-time was determined by fitting with an exponential one-phase association model (Y=Y0 + (Plateau – Y0)*(1-exp(-K*x))) where Y0 was set to zero and Plateau was set to one.

### Live-Cell Imaging

Live-cell and live-cell single-molecule imaging was performed on the same microscope described above. For live-cell imaging of HaloTagged DDR protein localization and DNA-damage induced foci, samples were densely labeled with 500 nm JF646 for 15 – 30 minutes, washed three times with complete medium and allowed to rest for 10 minutes before adding CO_2_-independent medium. For Zeocin-treated samples, cells were treated for one hour prior to HaloTag labeling with 100 µg/mL Zeocin. Z-stack images were acquired at 37ºC in the presence of 5% CO_2_ with a 100x objective and the 640 nm laser. These experiments were performed twice imaging >20 cells per experiment. For live-cell single-molecule imaging, HaloTagged DDR proteins were labeled with JFX650 at differing concentrations to achieve single-molecule density (0.1 nM for 30 seconds up to 20 nM for one minute). After labeling, cells were washed three times with complete medium and allowed to rest for 10 minutes before adding CO_2_-independent medium. For Zeocin-treated samples, cells were treated for one hour with Zeocin (100 µg/mL) prior to protein labeling with JFX650. Imaging was performed at 37ºC and 5% CO_2_ with the 100x objective and the 640 nm laser with highly inclined laminated optical sheet illumination (light angled between 1.29 and 1.32 depending on the sample). Images were acquired at 138 fps for 3000 frames followed by a brightfield image to visualize the cell.

### Analysis of Live-Cell Single-Molecule Imaging

Single-particle tracking (SPT) was performed in MATLAB 2019a using a version of SLIMfast allowing for analysis of TIFF files (Hansen et al., 2018). Settings for SPT used in the analysis were as follows: Exposure Time = 7.24 ms, NA = 1.49, Pixel Size = 0.16 µm, Emission Wavelength = 664 nm, *D*_max_ = 5 µm^2^/s, Number of gaps allowed = 2, Localization Error = -5, Deflation Loops = 0. While most HaloTagged DDR proteins exhibited near complete nuclear localization, REV7, SHLD3, SHLD2, and SHLD1 possessed mixed cytoplasmic and nuclear localizing fractions. For these proteins, the brightfield image was used to generate a nuclear mask in FIJI to separate cytoplasmic from nuclear tracks. Only nuclear tracks were used for analysis of protein diffusion in SpotOn. SPT files were then used in SpotOn in MATLAB to determine diffusion coefficients and the percentage of bound versus free particles. The following settings were used in SpotOn analysis of SPT files: TimeGap = 7.24 ms, dZ = 0.700 µm, GapsAllowed = 2, TimePoints = 8, JumpsToConsider = 4, BinWidth = 0.01 µm, PDF-fitting, D_Free_2State = [0.5 25], D_Bound_2State = [0.0001 0.5]. All live-cell single molecule imaging experiments were performed three times acquiring >20 cells per condition per experiment. Comparisons of Diffusion coefficients and fractions bound were performed in GraphPad Prism by two-way ANOVA with Tukey’s posthoc test.

## Supplemental Movie Legends

**Movie S1**. Representative movie demonstrating recruitment of JFX650-labeled 3xFLAG-HaloTagged DNA-PKcs to DNA DSBs after laser microirradiation. Images were acquired at one frame per second. 170×170 pixels with a pixel size of 0.16 µm.

**Movie S2**. Representative movie demonstrating recruitment of JFX650-labeled 3xFLAG-HaloTagged NBS1 to DNA DSBs after laser microirradiation. Images were acquired at one frame per second. 170×170 pixels with a pixel size of 0.16 µm.

**Movie S3**. Representative movie demonstrating recruitment of JFX650-labeled 3xFLAG-HaloTagged MDC1 to DNA DSBs after laser microirradiation. Images were acquired at one frame per second. 170×170 pixels with a pixel size of 0.16 µm.

**Movie S4**. Representative movie demonstrating recruitment of JFX650-labeled 3xFLAG-HaloTagged RNF168 to DNA DSBs after laser microirradiation. Images were acquired at one frame per second. 170×170 pixels with a pixel size of 0.16 µm.

**Movie S5**. Representative movie demonstrating recruitment of JFX650-labeled 3xFLAG-HaloTagged RNF169 to DNA DSBs after laser microirradiation. Images were acquired at one frame per second. 170×170 pixels with a pixel size of 0.16 µm.

**Movie S6**. Representative movie demonstrating recruitment of JFX650-labeled 3xFLAG-HaloTagged 53BP1 to DNA DSBs after laser microirradiation. Images were acquired at one frame every ten seconds. 170×170 pixels with a pixel size of 0.16 µm.

**Movie S7**. Representative movie demonstrating recruitment of JFX650-labeled 3xFLAG-HaloTagged RIF1 to DNA DSBs after laser microirradiation. Images were acquired at one frame every ten seconds. 170×170 pixels with a pixel size of 0.16 µm.

**Movie S8**. Representative movie demonstrating recruitment of JFX650-labeled 3xFLAG-HaloTagged SHLD3 to DNA DSBs after laser microirradiation. Images were acquired at one frame every ten seconds. 170×170 pixels with a pixel size of 0.16 µm.

**Movie S9**. Representative movie demonstrating recruitment of JFX650-labeled 3xFLAG-HaloTagged REV7 to DNA DSBs after laser microirradiation. Images were acquired at one frame every two seconds. 170×170 pixels with a pixel size of 0.16 µm.

**Movie S10**. Representative movie demonstrating recruitment of JFX650-labeled 3xFLAG-HaloTagged SHLD2 to DNA DSBs after laser microirradiation. Images were acquired at one frame every five seconds. 170×170 pixels with a pixel size of 0.16 µm.

**Movie S11**. Representative live-cell single-molecule imaging movies of untreated and zeocin-treated U2OS cells expressing 3xFLAG-HaloTagged NBS1 labeled with JFX650 and acquired at 138 frames per second. 170×140 pixels with a pixel size of 0.16 µm.

**Movie S12**. Representative live-cell single-molecule imaging movies of untreated and zeocin-treated U2OS cells expressing 3xFLAG-HaloTagged MDC1 labeled with JFX650 and acquired at 138 frames per second. 170×140 pixels with a pixel size of 0.16 µm.

**Movie S13**. Representative live-cell single-molecule imaging movies of untreated and zeocin-treated U2OS cells expressing 3xFLAG-HaloTagged RNF168 labeled with JFX650 and acquired at 138 frames per second. 170×140 pixels with a pixel size of 0.16 µm.

**Movie S14**. Representative live-cell single-molecule imaging movies of untreated and zeocin-treated U2OS cells expressing 3xFLAG-HaloTagged RNF169 labeled with JFX650 and acquired at 138 frames per second. 170×140 pixels with a pixel size of 0.16 µm.

**Movie S15**. Representative live-cell single-molecule imaging movies of untreated and zeocin-treated U2OS cells expressing 3xFLAG-HaloTagged 53BP1 labeled with JFX650 and acquired at 138 frames per second. 170×140 pixels with a pixel size of 0.16 µm.

**Movie S16**. Representative live-cell single-molecule imaging movies of untreated and zeocin-treated U2OS cells expressing 3xFLAG-HaloTagged RIF1 labeled with JFX650 and acquired at 138 frames per second. 170×140 pixels with a pixel size of 0.16 µm.

**Movie S17**. Representative live-cell single-molecule imaging movies of untreated and zeocin-treated U2OS cells expressing 3xFLAG-HaloTagged REV7 labeled with JFX650 and acquired at 138 frames per second. 170×140 pixels with a pixel size of 0.16 µm.

**Movie S18**. Representative live-cell single-molecule imaging movies of untreated and zeocin-treated U2OS cells expressing 3xFLAG-HaloTagged SHLD3 labeled with JFX650 and acquired at 138 frames per second. 170×140 pixels with a pixel size of 0.16 µm.

**Movie S19**. Representative live-cell single-molecule imaging movies of untreated and zeocin-treated U2OS cells expressing 3xFLAG-HaloTagged SHLD2 labeled with JFX650 and acquired at 138 frames per second. 170×140 pixels with a pixel size of 0.16 µm.

**Movie S20**. Representative live-cell single-molecule imaging movies of untreated and zeocin-treated U2OS cells expressing 3xFLAG-HaloTagged SHLD1 labeled with JFX650 and acquired at 138 frames per second. 170×140 pixels with a pixel size of 0.16 µm.

**Movie S21**. Representative live-cell single-molecule imaging movies of untreated and zeocin-treated U2OS cells expressing 3xFLAG-HaloTagged DNA-PKcs labeled with JFX650 and acquired at 138 frames per second. 170×140 pixels with a pixel size of 0.16 µm.

**Movie S22**. Representative live-cell single-molecule imaging movies of U2OS cells transiently expressing HaloTag-H2B and HaloTag-3xNLS labeled with JFX650 and acquired at 138 frames per second. 170×140 pixels with a pixel size of 0.16 µm.

**Movie S23**. Representative live-cell single-molecule imaging movies of HaloTagged MDC1 deletion mutants transiently expressed in ΔMDC1 U2OS cells, labeled with JFX650, and acquired at 138 frames per second. 170×140 pixels with a pixel size of 0.16 µm.

## Supplemental Figures

**Supplemental Figure 1.**
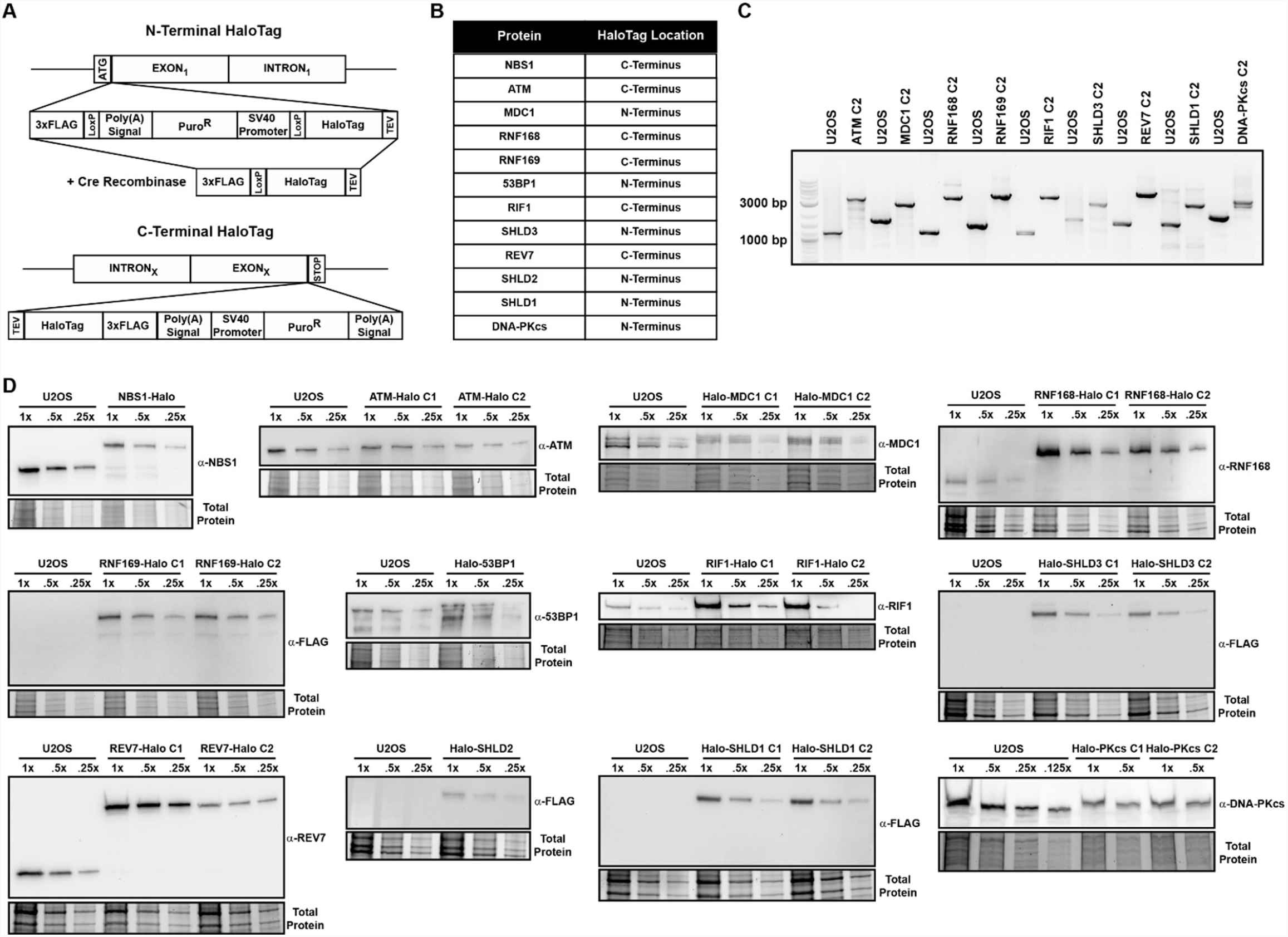
HaloTag knock-in design and knock-in validation. **A**. Graphical representation of the N-terminal and C-Terminal 3xFLAG-HaloTag used for genome editing. **B**. Table indicating the terminus where the HaloTag was introduced for each protein **C**. Agarose gel showing genomic PCR products and HaloTag insertion for second knock-in clones using primers oriented outside of the left and right homology arms **D**. Western blots for HaloTagged DDR proteins showing relative expression of each tagged protein compared to parental U2OS cells for proteins where a commercially available antibody was both available and capable of detecting endogenous expression levels. For proteins where protein-specific antibodies were not available, proteins were probed

**Supplemental Figure 2.**
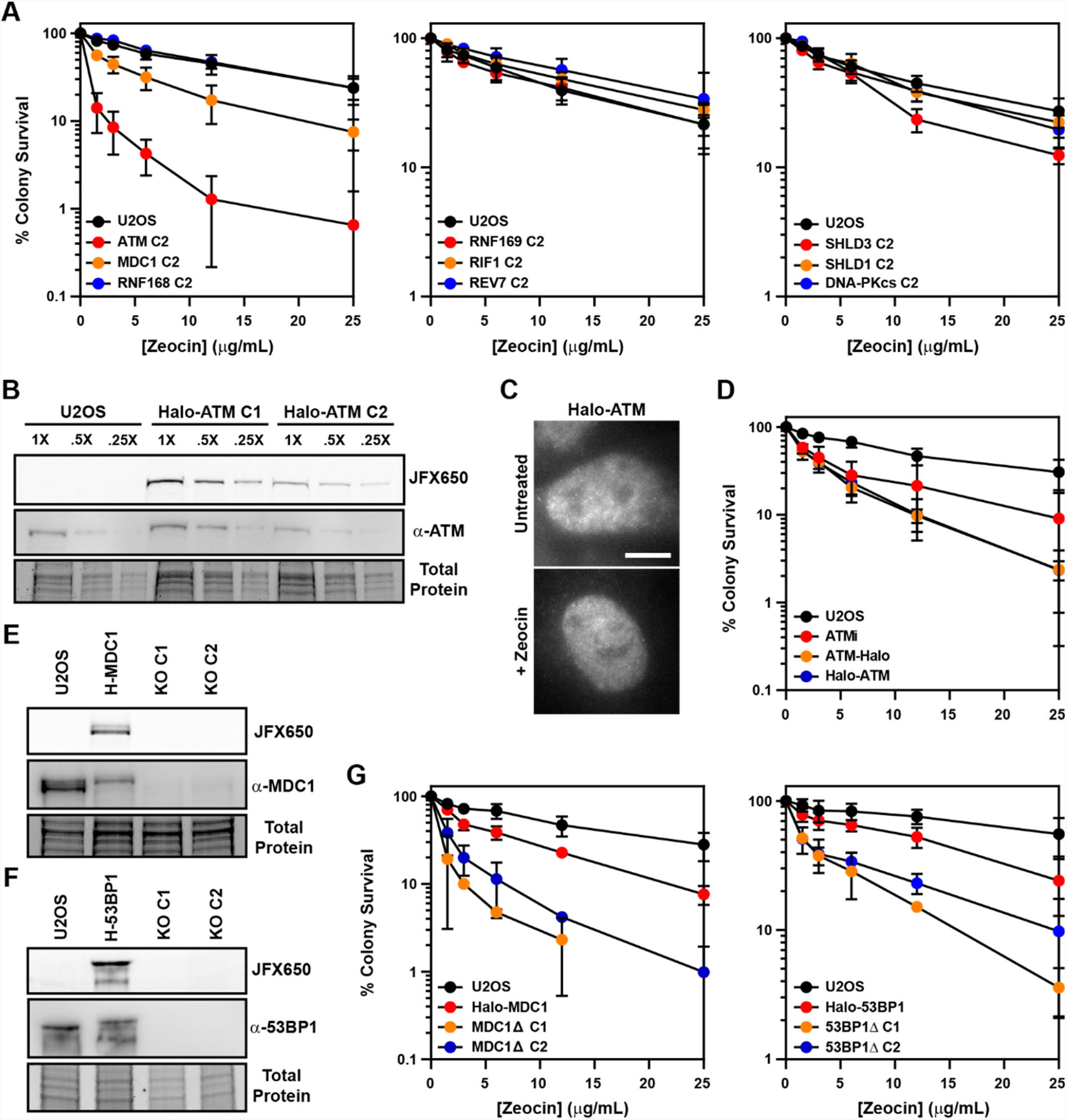
Functional validation of HaloTagged DDR proteins. **A**. Clonogenic survival assay showing sensitivity of second HaloTag clones to Zeocin relative to parental U2OS cells. **C**. SDS-PAGE and Western blot showing expression of N-terminally HaloTagged ATM. **C**. Representative image of live cells expressing Halo-ATM in the presence or absence of Zeocin. Scale bar = 10 µm. **D**. Clonogenic survival assays in the presence of absence of 10 µm ATMi (KU-55933) indicating that HaloTagging at either terminus abolishes ATM-mediated resistance to Zeocin. **E & F**. SDS-PAGE and Western blot showing absence of Halo-MDC1 (H-MDC1) or Halo-53BP1 (H-53BP1) expression in MDC1 and 53BP1 knockout cells. **G**. Clonogenic survival assays of MDC1 and 53BP1 knockout clones showing sensitivity to Zeocin relative to Halo-MDC1 or Halo-53BP1 and U2OS cells. Clonogenic survival assays are presented as the average ± S.D. of at least 3 independent

**Supplemental Figure 3.**
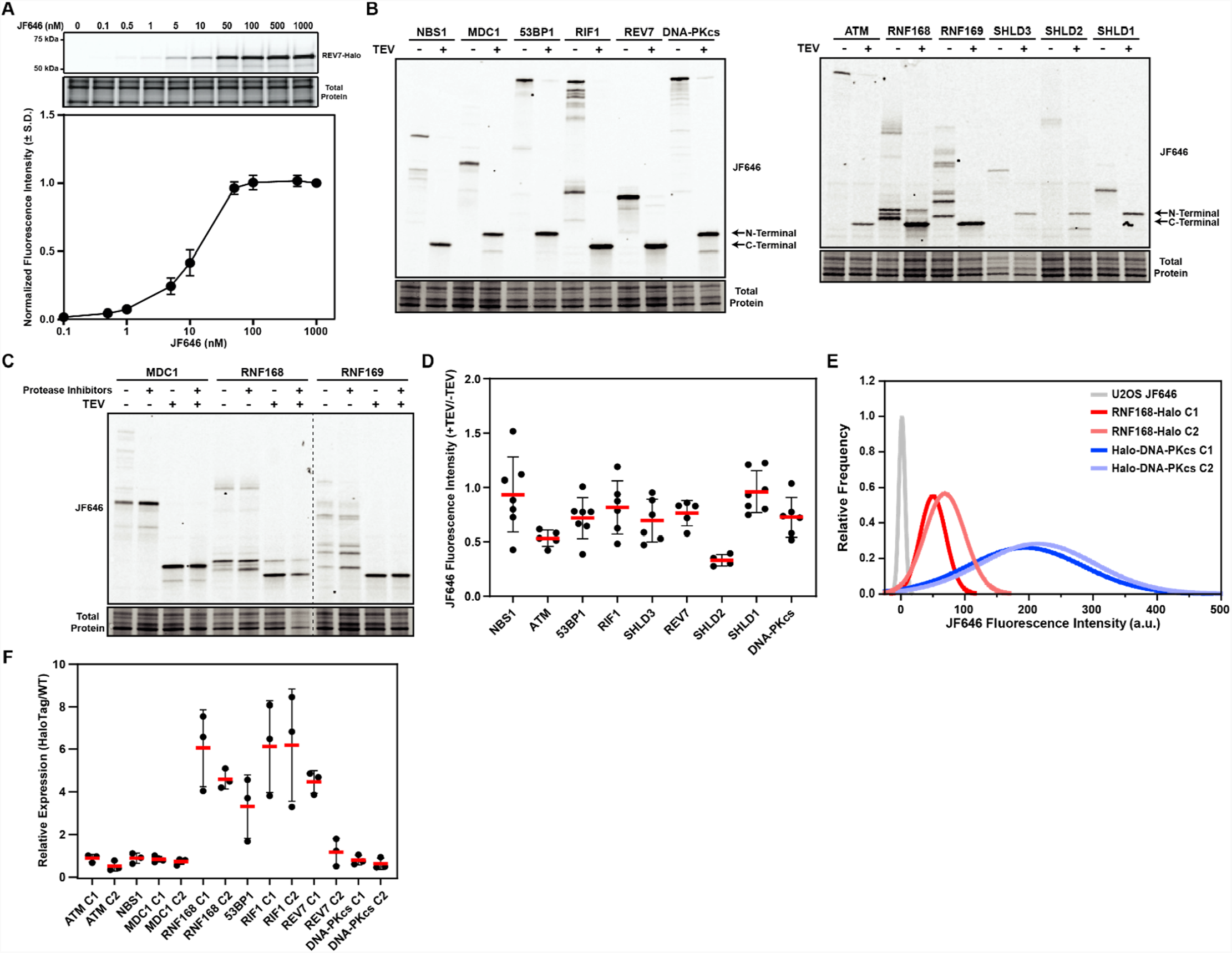
Quantification of absolute protein abundance using HaloTagged DDR proteins. **A. HaloTag labeling of** REV7-Halo in cells titrated with increasing concentrations of JF646 with an associated saturation curve. Saturation curve data is plotted as the mean ± S.D. from three independent experiments. **B**. Representative in-gel fluorescence images for TEV-digestion experiments used to generate an adjustment factor for protein abundance based upon JF646 signal intensity. N-terminal refers to the N-terminal 3XFLAG-HaloTag and C-Terminal refers to the C-terminal 3XFLAG-HaloTag. **C**. In-gel fluorescence showing protein degradation products of MDC1, RNF168, and RNF169 in the presence or absence of protease inhibitor cocktail after lysing in CHAPS lysis buffer. **D**. Quantification of TEV digestion experiments generating a TEV adjustment factor for absolute protein abundance. Each data point represents an independent experiment. Red bar indicates the mean and error bars represent S.D. **E**. Representative plots of JF646 fluorescence intensity in parental U20S cells, two RNF168-Halo clones, and two Halo-DNA-PKcs clones detected by flow cytometry. **F**. Quantification of expression of HaloTagged protein relative to untagged protein in U20S cells to generate a correction factor which was used to calculate the number of molecules per cell for each protein in wild-type U20S cells. Data are quantified from three independent experiments. Red bar indicates the mean and error bars represent S.D.

**Supplemental Figure 4.**
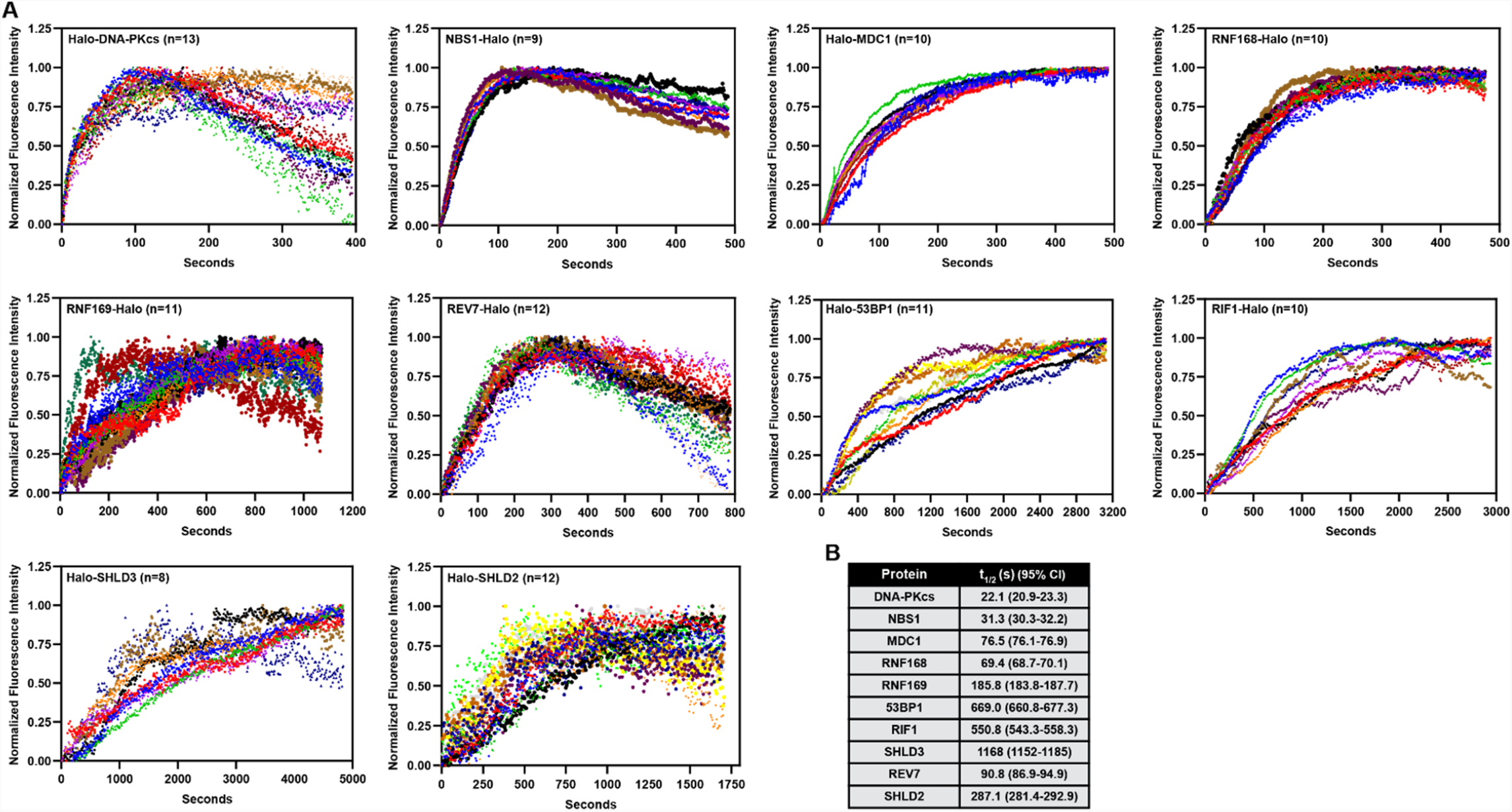
Kinetics of HaloTagged DDR protein recruitment to laser-induced DSBs. **A**. Plots showing normalized fluorescence intensity and recruitment of each HaloTagged DDR protein to laser-induced DSBs plotted by individual cells rather than normalized average. Each color represents an individual cell that was analyzed. Normalized fluorescence intensity was done by setting the brightest frame for each cell to 1. **B**. Table listing the recruitment half-times (t1/2) of each protein to laser-induced DSBs determined by nonlinear regression assuming one-phase association.

**Supplemental Figure 5.**
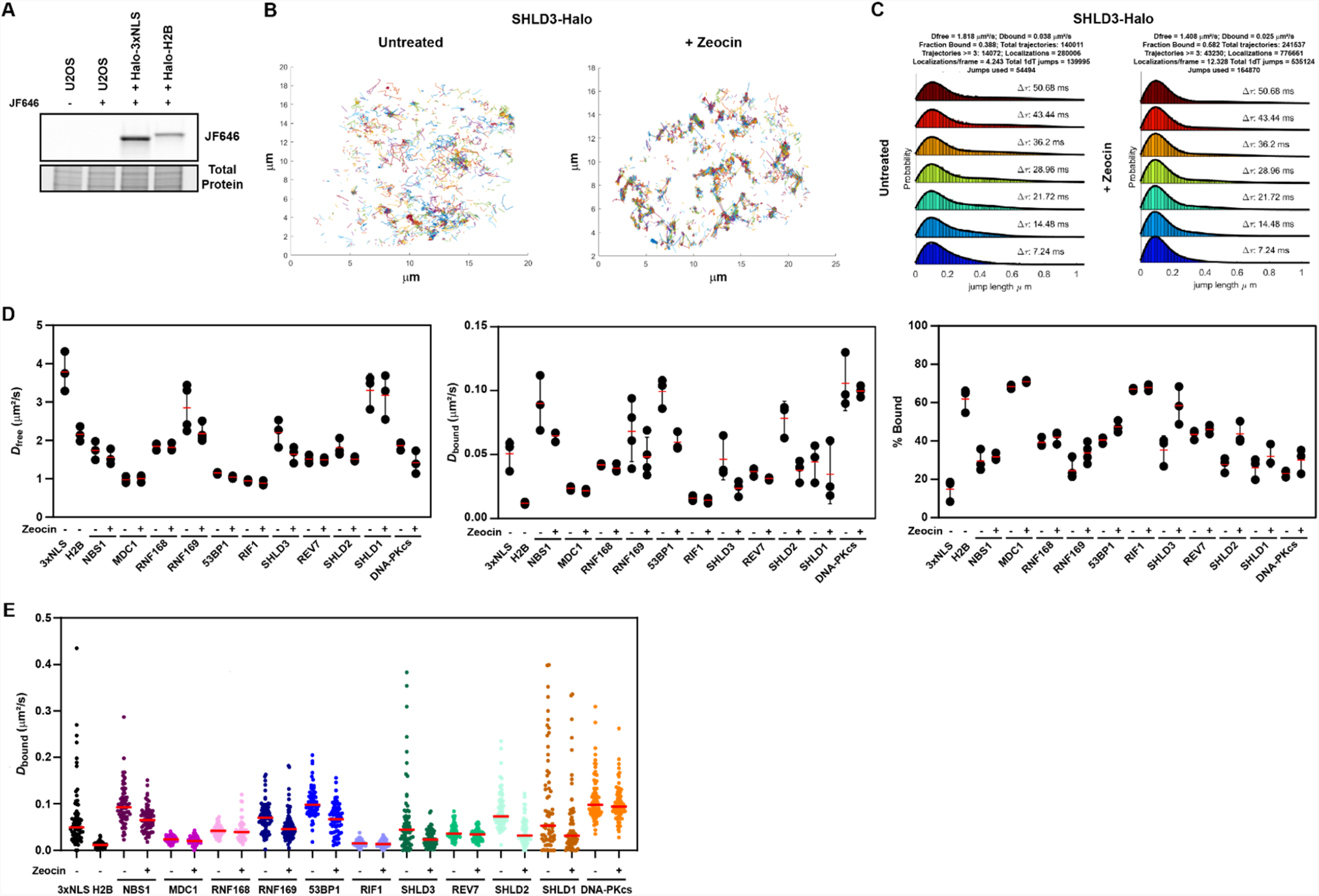
Nuclear diffusion and chromatin binding characteristics of HaloTagged DDR proteins. A. SDS-PAGE gel showing expression of Halo-H2B and Halo-3xNLS in cells labeled with JF646. **B**. Example images of protein tracks generated by single particle tracking of Halo-SHLD3 ± Zeocin. **C**. Example plots from SpotOn for a single live-cell single-molecule experiment (>20 cells) for Halo-SHLD3 ± Zeocin. **D**. Same data presented in Figure 5B & C, except plotted by individual experiment, rather than as individual cells, to demonstrate the reproducibility of imaging between experiments. Each dot represents results from an individual experiment. Red bar = mean. Error bars = S.D. **E**. Diffusion coefficients of the bound fraction of HaloTag DDR proteins present in at least 3 consecutive frames in untreated conditions and post-Zeocin exposure. Values plotted indicate the *D*_bound_ for all analyzed tracks per cell with each dot indicating a separate cell that was analyzed. Live-cell single-molecule imaging was performed over 3-4 separate days imaging at least 20 individual cells per condition per experimental replicate (n ≥ 60 cells total for each protein and condition).

**Supplemental Figure 6.**
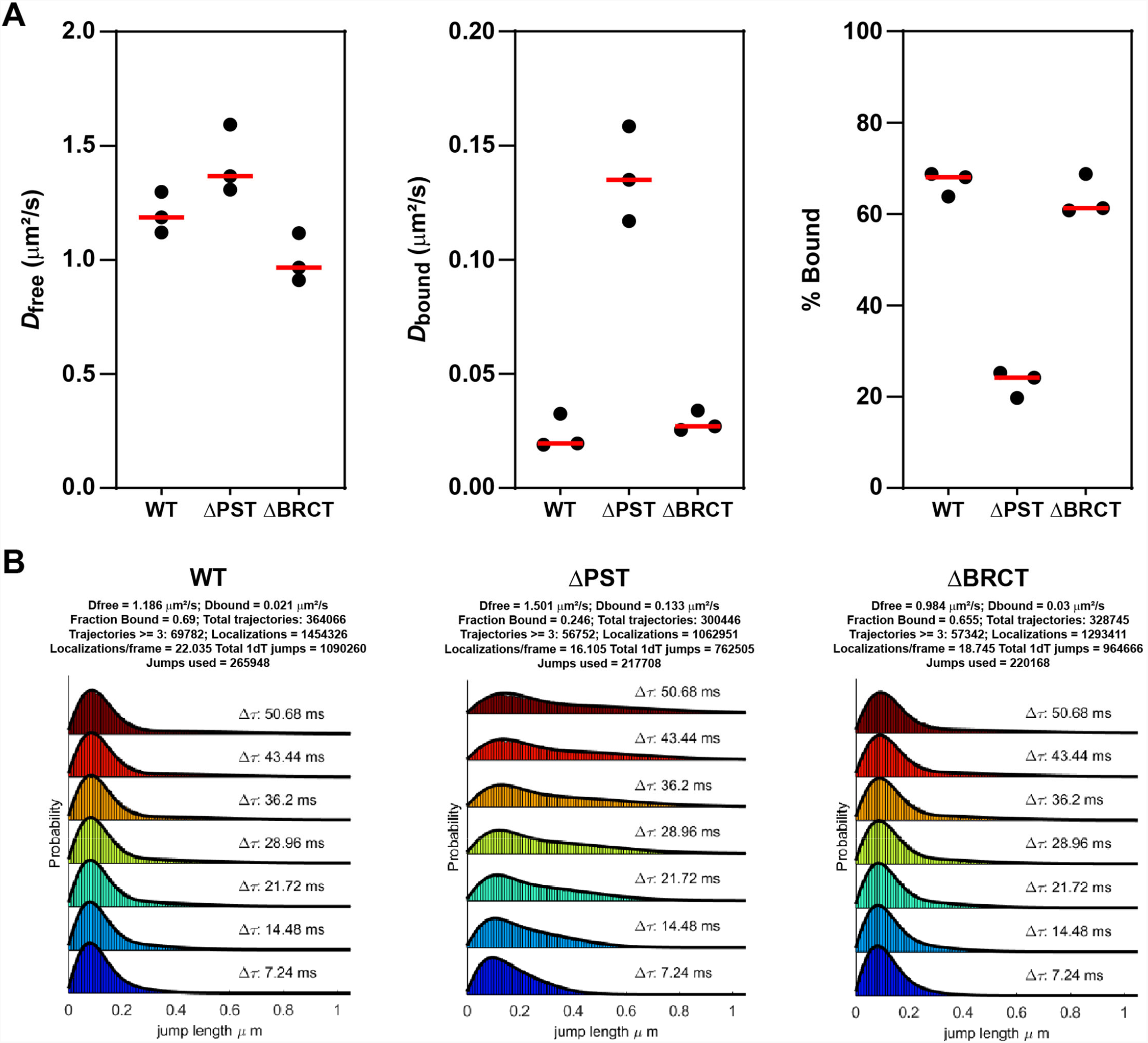
**A.** Same data presented in Figure 6D, except plotted by individual experiment, rather than individual cells, demonstrating the reproducibility of analysis experiments. Each dot represents results from an individual experiment. Red bar = median. **B**. Example plots generated by SpotOn for a single live-cell single-molecule experiment (>20 cells) of WT, ΔPST, and ΔBRCT MDC1.

